# Lanthanide-dependent isolation of phyllosphere methylotrophs selects for a phylogenetically conserved but metabolically diverse community

**DOI:** 10.1101/2023.06.28.546956

**Authors:** Alekhya M. Govindaraju, Colleen A. Friel, Nathan M. Good, Sidney L. Banks, Kenan S. Wayne, N. Cecilia Martinez-Gomez

## Abstract

Lanthanides have emerged as important metal cofactors for biological processes. Lanthanide-associated metabolisms are well-studied in leaf symbiont methylotrophic bacteria, which utilize reduced one-carbon compounds such as methanol for growth. Yet, the importance of lanthanides in plant-microbe interactions and on microbial physiology and colonization in plants remains poorly understood. To investigate this, 344 pink-pigmented facultative methylotrophs were isolated from soybean leaves by selecting for bacteria capable of methanol oxidation with lanthanide cofactors, but none were obligately lanthanide-dependent. Phylogenetic analyses revealed that all strains were nearly identical to each other and are part of the *extorquens* clade of *Methylobacterium*, despite variability in genome and plasmid sizes. Strain-specific identification was enabled by the higher resolution provided with *rpoB* compared to 16S rRNA as marker genes. Despite the low strain-level diversity, the metabolic capabilities of the collection diverged greatly. Strains encoding identical lanthanide-dependent alcohol dehydrogenases displayed significantly different growth rates and/or final ODs from each other on alcohols in the presence and absence of lanthanides. Several strains also lacked well-characterized lanthanide-associated genes thought to be important for phyllosphere colonization. Additionally, 3% of our isolates were capable of growth on sugars and 23% were capable of growth on aromatic acids, substantially expanding the range of substrates utilized by *Methylobacterium extorquens* in the phyllosphere. Our findings suggest that the expansion of metabolic capabilities, as well as differential usage of lanthanides and their influence on metabolism, among closely related strains point to evolution of niche partitioning strategies to promote colonization of the phyllosphere.

**Importance:** Lanthanide metals have long been appreciated for their role in technology, but the recent identification of lanthanides as cofactors in methylotrophic metabolism has expanded the role of lanthanides into biology. In the phyllosphere, methylotrophs are some of the most abundant bacteria found on leaf surfaces, where lanthanide concentrations are sufficiently high to support their growth. Yet, the extent to which lanthanides influence methylotrophic metabolism in the phyllosphere remains unknown. Here, we characterize a methylotrophic enrichment isolated from the phyllosphere in a lanthanide-dependent manner. We have identified (1) closely related strains with identical lanthanide-dependent enzymes that exhibit different growth on alcohols in the presence of lanthanides, (2) resilient strains lacking lanthanide-associated genes thought to be important for phyllosphere colonization, and (3) many strains capable of metabolisms that were thought to be rare within this clade. Overall, our isolates serve as a model community to interrogate how lanthanides differentially influence methylotrophic physiology.

## Introduction

The phyllosphere, the aerial portion of plants dominated by leaves (1,2), is colonized by a diverse community of microorganisms with populations approaching 10^7^ bacteria per square cm of leaf (2,3). Leaf surfaces can provide a variety of substrates to support bacteria, including carbohydrates, organic acids, amino acids, and volatile compounds(3). Yet, bacteria in the phyllosphere must overcome waxy leaf cuticles, UV light, temperature fluctuations, low water availability, and other biotic and abiotic impediments to colonization (1–4). Heterogeneous leaf microenvironments thus require strains with high resource overlap to effectively niche partition to balance coexistence with competition (5).

A study characterizing the phyllosphere community of four phylogenetically distinct plants – *Glycince max* (soybean), *Trifolium repens* (white clover), *Arabidopsis thaliana*, and *Oryza sativa* (rice) —found that all four plant species showed consistently high relative abundance of the genus *Methylobacterium* (6). Methylotrophs of the *Methylobacterium* genus are so-named due to their capacity to utilize reduced one-carbon compounds such as methanol, methylamine, formaldehyde, or formate as their sole source of carbon and energy (7,8). In the phyllosphere, methylotrophs are well-suited to take advantage of methanol released daily by pectin methylesterases during routine plant cell wall breakdown and repair (3,7,9).

Methylotrophs may even stimulate plants into releasing additional methanol by secreting the plant hormone cytokinin, which triggers plant cell division (2,7,10). *Methylobacterium* species can be further classified as pink-pigmented facultative methylotrophs (PPFMs) that can utilize multi-carbon substrates available in the phyllosphere (7,8,11). This includes organic acids such as acetate (C2), pyruvate (C3), and succinate (C4) that are assimilated via the TCA cycle (8) and methoxylated aromatic acids that can broken down into intermediates that can feed into both methylotrophic pathways and the TCA cycle (12). The capability to utilize different carbon substrates, may explain why methylotrophs are so well-suited to life in the phyllosphere (1,3,10,11) where metabolite availability changes dynamically (7).

Metagenomic and metaproteomic studies of the phyllosphere have consistently detected high abundance of proteins involved in one-carbon metabolism (6,13,14), specifically, the pyrroloquinoline quinone (PQQ) methanol dehydrogenases, MxaFI and XoxF1, that convert methanol to formaldehyde. The calcium-dependent MxaFI was the first-identified methanol dehydrogenase from Gram-negative methylotrophs and thought to be the sole enzyme responsible for methanol oxidation in this system (15–17). However, recent studies have highlighted the importance of an alternative methanol dehydrogenase, XoxF1, which coordinates a lanthanide atom in complex with PQQ (18). XoxF1 is widespread in the environment (19,20) and phylogenetically ancestral to MxaFI (21). Studies investigating both methanol dehydrogenase systems have shown that induction of *xoxF* can repress *mxaFI* expression at lanthanide concentrations higher than 100 nM (22). Additional work has led to the identification of ExaF (23), a lanthanide-dependent alcohol dehydrogenase with sub-nanomolar *K*_M_ towards ethanol and auxiliary activity with formaldehyde (24), a toxic intermediate of methylotrophic metabolism.

*Methylobacterium extorquens* AM1 (**<u>A</u>**irborne **<u>M</u>**ethylotroph #**<u>1</u>**) has been a model for studying aerobic facultative methylotrophy for decades due to its genetic tractability and reproducible growth phenotypes (25–27). Yet, its long history of laboratory domestication and large number of insertion sequences (28) has led researchers to seek out other *M. extorquens* strains with comparable metabolisms and genetic tractability, such as *Methylobacterium extorquens* PA1 (**<u>P</u>**hyllosphere **<u>A</u>**rabidopsis #**<u>1</u>**). *M. extorquens* PA1 shares 100% 16S rRNA identity with *M. extorquens* AM1, has fewer insertion sequences, and has ecological relevance (4).

*M. extorquens* AM1 and PA1 have emerged as models for understanding the role of lanthanides in methylotrophy. Lanthanide concentrations in the phyllosphere of *Arabidopsis thaliana* have been measured to be 7-10 μg per g plant dry weight (29), which is sufficient to drive lanthanide-dependent microbial processes in *M. extorquens* (22). Additionally, lanthanides have been shown to have positive and/or hormetic effects on plants themselves (30,31). Despite extensive characterizations of the microbial community in the phyllosphere as well as recent advances in lanthanide-dependent metabolisms, the effect of lanthanides on methylotrophic bacterial composition in the phyllosphere remains unknown. To investigate this, methylotrophs were isolated from the phyllosphere of soybean plants and selected for growth on methanol in the presence and absence of lanthanides, as well as growth on additional substrates. Growth parameters were compared to those of *M. extorquens* AM1 – a model organism for studying lanthanide-dependent methylotrophy (18,22,24,32)– and *M. extorquens* PA1 – a model plant commensal organism (4,29). As compared to other members of the *extorquens* clade, our isolated methylotrophic strains were found to be phylogenetically similar but phenotypically distinct, with expanded metabolic capabilities and growth rates differentially affected by substrate identity and the presence or absence of lanthanum.

## Results

### The soybean phyllosphere methylotrophic community is not obligately lanthanide-dependent

Methylotrophic bacterial strains were isolated from the soybean phyllosphere by selection on minimal PIPES-buffered media (33) with methanol as the sole carbon source. This medium has been optimized for robust and reproducible growth of diverse *Methylobacterium* strains (33). Previous studies have shown that the presence of lanthanides allows for the isolation of novel methylotrophic strains and/or metabolic capabilities (34–37); thus, we also included lanthanum in our media. Of 344 total isolates, 158 strains were isolated with the addition of La^3+^ and 186 strains without the addition of La^3+^ to the selection medium. After isolating single colonies, all 344 strains were retested for growth on methanol in the presence and absence of La^3+^ to determine if lanthanides were necessary for growth. All strains were pink, showed growth on methanol in the presence and absence of La^3+^ in the growth medium, and showed growth on succinate and were therefore all classified as PPFMs. Each strain was assigned a unique SLI (**S**oybean **L**eaf **I**solate) number, which is used to reference specific strains throughout this study (**Table 1**).

**Table 1.** Summary of SLI strain genomes. Whole-genome sequencing performed by Joint Genome Institute using IMG Annotation Pipeline v.5.0.22. Chromosome, (C); Plasmid (P).

| Strain | IMG ID | Size (bp)<br>(C) | % GC<br>(C) | Genes<br>(C) | Size (bp)<br>(P) | % GC (P) | Genes<br>(P) |
| --- | --- | --- | --- | --- | --- | --- | --- |
| *M. extorquens*<br>AM1 | 644736386 | 5511322 | 68.71 | 5029 | 1261460 | 67.65 | 1168 |
|  |  |  |  |  | 44195 | 67.93 | 33 |
|  |  |  |  |  | 37858 | 65.25 | 34 |
|  |  |  |  |  | 24943 | 66.94 | 30 |
| *M. extorquens*<br>PA1 | 641228497 | 5471154 | 68.18 | 4939 | - | - | - |
| *M. sp.* AMS5 | 2684623050 | 5435450 | 68.50 | 4970 | 117697 | 65.29 | 162 |
|  |  |  |  |  | 25608 | 65.30 | 24 |
|  |  |  |  |  | 20451 | 66.33 | 27 |
| *M. extorquens*<br>SLI 223 | 2917523761 | 5504362 | 68.23 | 5222 | - | - | - |
| *M. extorquens*<br>SLI 231 | 2918956041 | 5373034 | 68.29 | 5165 | 166358 | 67.47 | 176 |
|  |  |  |  |  | 27292 | 61.51 | 25 |
|  |  |  |  |  | 29644 | 58.78 | 27 |
| *M. extorquens*<br>SLI 233 | 2932223180 | 5373474 | 68.29 | 5287 | 166399 | 67.47 | 181 |
|  |  |  |  |  | 27296 | 61.51 | 28 |
|  |  |  |  |  | 29645 | 58.79 | 27 |
| *M. extorquens*<br>SLI 274 | 2947147188 | 5374052 | 68.29 | 5202 | 166386 | 67.47 | 178 |
|  |  |  |  |  | 27292 | 61.51 | 26 |
|  |  |  |  |  | 29646 | 58.79 | 27 |
| *M. extorquens*<br>SLI 285 | 2932228704 | 5375017 | 68.30 | 5295 | 166399 | 67.47 | 182 |
|  |  |  |  |  | 27292 | 61.51 | 25 |
|  |  |  |  |  | 29655 | 58.78 | 28 |
| *M. extorquens*<br>SLI 384 | 2918961435 | 5504365 | 68.23 | 5233 | 28462 | 67.45 | 48 |
| *M. extorquens*<br>SLI 499 | 2947152626 | 5373409 | 68.29 | 5218 | 166383 | 67.47 | 178 |
|  |  |  |  |  | 27292 | 61.51 | 25 |
|  |  |  |  |  | 29645 | 58.79 | 27 |
| *M. extorquens*<br>SLI 505 | 2918966707 | 5372985 | 68.29 | 5170 | 166379 | 67.47 | 176 |
|  |  |  |  |  | 27292 | 61.51 | 26 |
|  |  |  |  |  | 29645 | 58.79 | 27 |
| *M. sp.* SLI 516 | 2918972107 | 5009907 | 68.66 | 4762 | 199519 | 67.26 | 215 |
|  |  |  |  |  | 28884 | 66.85 | 39 |
|  |  |  |  |  | 20194 | 60.93 | 30 |
| *M. p.* SLI 575 | 2918977154 | 5009893 | 68.66 | 4764 | 199519 | 67.26 | 214 |
|  |  |  |  |  | 28890 | 66.84 | 39 |
|  |  |  |  |  | 20194 | 60.93 | 30 |
| *M. sp.* SLI 576 | 2918982202 | 5009856 | 68.66 | 4762 | 199518 | 67.26 | 215 |
|  |  |  |  |  | 28890 | 66.84 | 39 |
|  |  |  |  |  | 20194 | 60.93 | 31 |

### Methylotrophic communities from the soybean phyllosphere are phylogenetically similar

Chromosomal DNA was extracted from the isolates and *rpoB*, the gene that encodes for the beta subunit of RNA polymerase, was PCR amplified and Sanger sequenced. Using NCBI BLAST, *rpoB* sequences from each isolate were compared against the Joint Genome Institute’s (JGI) Integrated Microbial Genomes (IMG) database to putatively assign species-level taxonomic classifications to each isolate. All isolates matched to *Methylobacterium* species within group B (38), with the closest species being *Methylobacterium extorquens* AM1, *Methylobacterium extorquens* PA1, *Methylobacterium extorquens* TK001, and *Methylobacterium zatmanii* 135. Untrimmed *rpoB* sequences from each isolate, as well as from reference *Methylobacterium* and related species, were aligned using MUSCLE, and Maximum-Likelihood trees with 100 bootstrap replications were constructed using MEGA (**Figure 1A**). The *rpoB*-based trees show a high degree of relatedness amongst all isolates, integrating SLI strains among other reference methylotroph strains within the *extorquens* clade.

**Figure 1.**
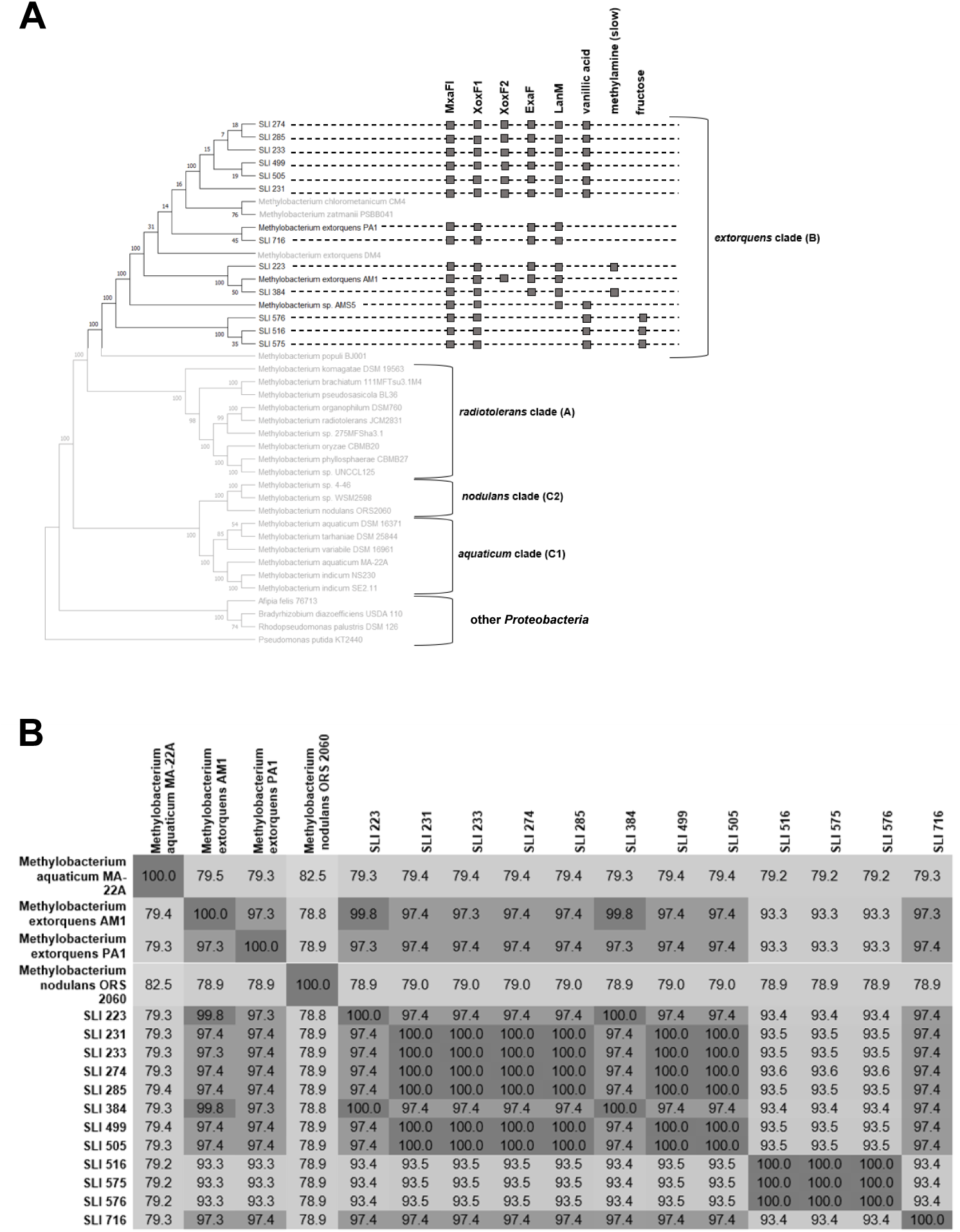
Phylogenetic and taxonomic characterization of a methylotrophic community isolated from the soybean phyllosphere. (**A**) Phylogenetic tree based on *rpoB* sequences. Untrimmed *rpoB* sequences from each isolate as well as from reference *Methylobacterium* and related species were obtained from IMG, aligned using MUSCLE, and the Maximum-Likelihood trees were constructed with 100 bootstrap replications using MEGA. Groupings indicate clade identity for each strain. Boxes indicate metabolic capabilities (growth on vanillic acid, growth on fructose, slow growth on methylamine) and lanthanide-related gene products (MxaFI, XoxF, ExaF, LanM) present in SLI strains and model *extorquens* clade species referenced in this study (*M. extorquens* AM1, *M. extorquens* PA1, *M. sp.* AMS5). (**B**) Average nucleotide identity of all SLI strains compared to representative species from the *extorquens, nodulans,* and *aquaticum* clades. All pairwise comparison ANIs found in **Table S1**

The genomes of SLI strains 223, 231, 233, 274, 285, 384, 499, 505, 516, 575, 576 were sequenced through the Joint Genome Institute and annotated using the IMG Annotation Pipeline v.5.0.22. We sought to augment our understanding of the SLI collection composition at the taxonomic level by employing pairwise average nucleotide identity (ANI) comparisons of all orthologous genes shared among all assembled SLI genomes (see **Table 1**) and representative methylotrophic genomes using the IMG database. Results from the complete pairwise ANI comparisons can be found in the **Table S1**, but have been condensed to include only SLI strains and representative species from the *extorquens, aquaticum,* and *nodulans* clades in **Figure 1B**. Based on ANI comparisons, all SLI strains were most similar to *Methylobacterium* species from the *extorquens* clade (93-99% ANI) when compared to the *nodulans* (79% ANI) and *aquaticum* clades (78% ANI), indicating that our collection likely belongs to the *extorquens* clade. Based on the 95% standard species cutoff (39), all SLI strains except for SLI 516, 575, and 576 can be classified as *Methylobacterium extorquens* strains when compared to *M. extorquens* AM1 and *M. extorquens* PA1. SLI 516, 575, and 576 cluster within the *extorquens* clade but have ANI values under 95%; thus, these isolates are not of the *extorquens* species and the presence of more than one species confirms that our isolates are a community. Additionally, SLI 231, 233, 274, 285, 499, and 505 are genetically identical to each other; yet, as will be discussed in detail later, they exhibit distinct metabolic differences (see **Table S2** for full comparison of genomes).

### Whole-genome sequencing of representative SLI strains reveals new *extorquens* strains

Summaries of the genome information for each sequenced SLI strain compared to *M. extorquens* AM1 and PA1 are represented in **Table 1**. Notably, all strains but SLI 223 encoded at least one plasmid in addition to the chromosome, with most strains (SLI 231, 233, 274, 285, 499, 505, 516, 575, 576) encoding at least three additional plasmids. The size, % GC, and number of genes per chromosome and per plasmid were very similar for SLI 231, 233, 274, 285, 499, and 505 and for SLI 516, 575, 576; this underscores the high degree of ANI between the strains (**Figure 1B)**. SLI 516, 575, and 576 – which are the least similar by ANI to *M. extorquens* AM1 and PA1 (**Figure 1B**) – also have the smallest chromosome sizes despite having the broadest metabolic capabilities (**Figure 2A**). None of the strains encoded the megaplasmid (1261460 bp) found in *M. extorquens* AM1, and the plasmid sizes of the SLI strains were very different from the plasmid sizes of *M. extorquens* AM1.

**Figure 2.**
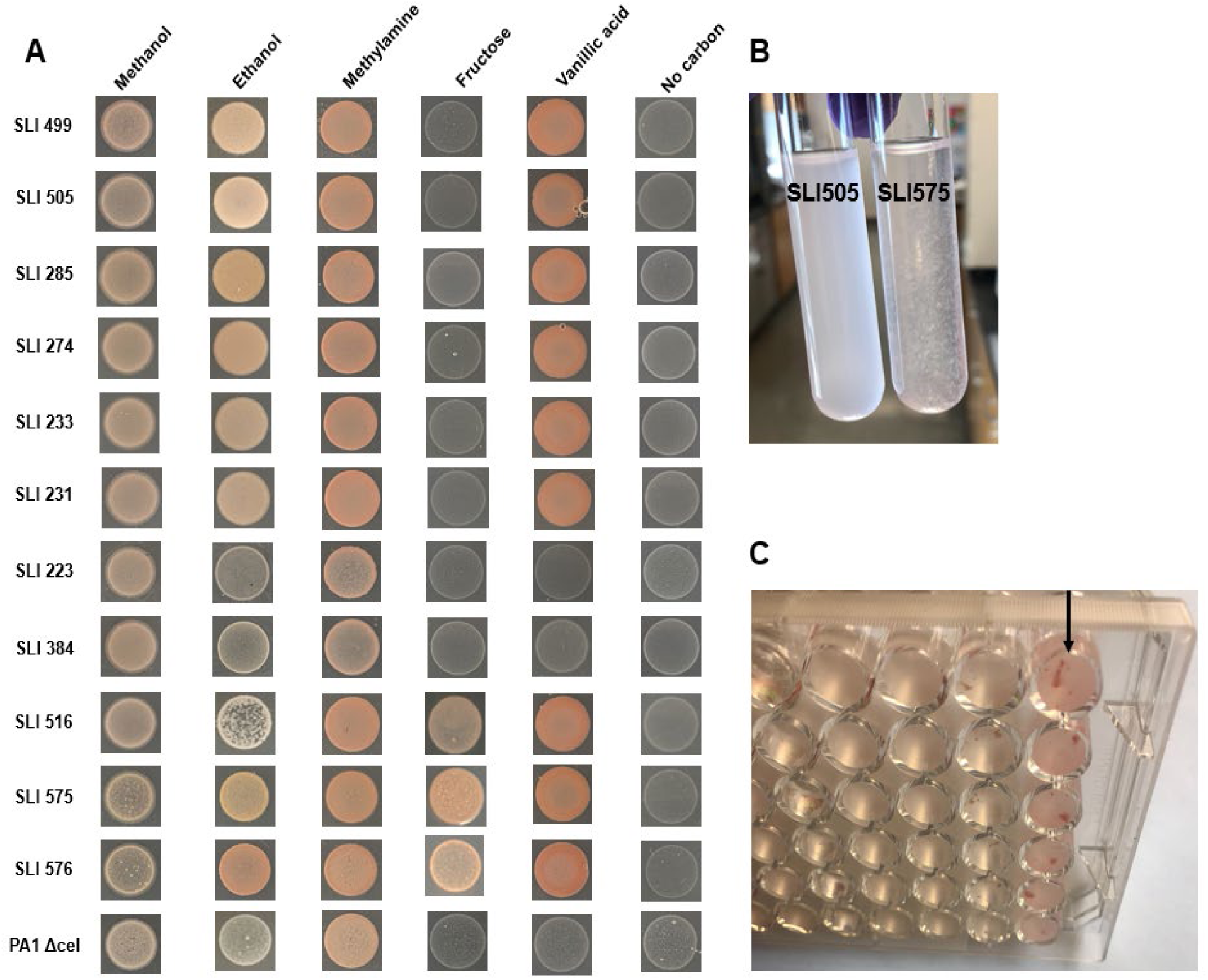
**A.** Spotting of a subset of SLI strains on various substrates to demonstrate differential metabolic capabilities of SLI test cohort. All strains were spotted on minimal agar supplemented with either 20 mM methanol, 34 mM methanol, 15 mM methylamine, 5 mM vanillic acid, 25 mM fructose, or no carbon. *Methylobacterium extorquens* PA1Δ*cel* as a comparison of known substrate utilization capabilities in a model *extorquens* clade strain. Biofilm formation (verified by crystal violet staining, data not shown) of SLI 575 in liquid culture **(B**) and in microplates (**C**)

Predictions about metabolic potential can be made from genomic analysis (40) (see **Table S2**). All SLI genomes encode tetrahydromethanopterin and tetrahydrofolate-dependent pathways for C_1_ oxidation and marker genes for the serine cycle for C_1_ assimilation, indicating that all strains are Type II methylotrophs (8,27,40,41). All SLI genomes encode four formate dehydrogenases, save for SLI 516, 575, and 576 which lack formate dehydrogenase 1. All SLI genomes also encode the *N*-methylglutamate pathway for methylamine utilization (*mgdDCBA*, *mgsABC*, *gmaS*) (42). Strains SLI 231, 233, 274, 285, 499, 505, 516, 575, and 576 encode a previously identified gene island that confers the ability to grow on methoxylated aromatic acids (12).

The SLI genomes can be grouped into three categories based on the presence of certain lanthanide-associated genes: group 1 consists of SLI 231, 233, 274, 285, 499, 505; group 2 consists of SLI 223 and SLI 384; group 3 consists of SLI 516, 575, and 576. All genomes encode calcium-dependent methanol dehydrogenases (*mxaFI*) and at least one lanthanide-dependent methanol dehydrogenase (*xoxF1)* indicating that all strains are capable of lanthanide-independent and lanthanide-dependent methanol metabolism. Additionally, all strains encode genes for the TonB-dependent and ABC transporters of the lanthanide utilization and transport (*lut)* cluster (43) and encode genes with high degrees of similarity to the biosynthetic gene cluster for the lanthanide chelator, methylolanthanin, in *M. extorquens* AM1 (44). Yet, the gene for the XoxF1 paralog, XoxF2 (22), is absent in genomes from groups 2 and 3 and the gene for the lanthanide-dependent alcohol dehydrogenase (*exaF*) is lacking in genomes from group 3. In addition, group 3 genomes lack the gene for the lanthanide-binding protein, lanmodulin (*lanM*), despite encoding the rest of the genes involved in the lanthanide utilization and transport cluster (43,45). For comparison, *M. extorquens* AM1 encodes MxaFI, XoxF1, XoxF2, ExaF, and LanM similar to SLI strains from group 1 and *M. extorquens* PA1 encodes MxaFI, XoxF1, ExaF, and LanM similar to SLI strains from group 2. Thus, the influence of lanthanides on the metabolisms of group 3 strains is of particular interest as they only encode a single lanthanide-associate protein, XoxF1. The differences between groups 1, 2, and 3 is also reflected in the average nucleotide identities in **Figure 1B**. A complete summary of all metabolic and lanthanide-related genes found in the SLI strains is included in **Table S2**.

### Methylotroph isolates possess broad metabolic capabilities independent of lanthanide availability

Although phyllosphere isolates were phylogenetically similar to domesticated research strains such as *M. extorquens* AM1 and PA1, we hypothesized that environmental methylotrophs might possess expanded substrate repertoires and broader metabolic capabilities (4,5). To test our hypothesis, all isolates were screened for growth on C_1_ substrates (methanol, methylamine, dimethylsulfide), organic acids (oxalate, succinate), other alcohols (ethanol), sugars (fructose, glucose), aromatic acids (vanillic acid), and complex insoluble substrates (lignin, cellulose, tyrosine) **(Table 2)**. Lanthanides were also added to growth media for each substrate, but this addition did not unlock novel substrate utilization capabilities for any SLI tested. All strains exhibited growth on methanol, methylamine, succinate, and ethanol. No strains grew on dimethylsulfide, oxalate, glucose, lignin, cellulose, or tyrosine. Notably, 10 out of 344 strains grew on fructose and 78 out of 344 strains grew on vanillic acid.

**Table 2.** Distribution of substrate-utilization capabilities of SLI community. All 344 isolates were grown in liquid minimal salts media containing either C_1_ (20 mM methanol, 15 mM methylamine, 1 mM dimethylsulfide), C_2_ (34 mM ethanol, 5 mM potassium oxalate), C_4_ (succinate), sugars (25 mM fructose, 25 mM glucose), or complex insoluble substrates (vanillic acid, cellulose, lignin, tyrosine)

| **Substrate** | **No. of SLI strains** |
| --- | --- |
| Methanol | 344 |
| Methylamine | 344 |
| Dimethylsulfide | 0 |
| Ethanol | 344 |
| Potassium Oxalate | 0 |
| Succinate | 344 |
| Glucose | 0 |
| Fructose | 10 |
| Vanillic Acid | 78 |
| Lignin | 0 |
| Cellulose | 0 |
| Tyrosine | 0 |

The subset of SLI strains with completely sequenced genomes (SLI 223, 231, 233, 274, 285, 384, 499, 505, 516, 575, 576) that were shown to have broad metabolic capabilities from the screen described above were spotted onto solid media containing five different carbon sources without La^3+^, with growth of *M. extorquens* PA1 Δ*cel* (a mutant shown to reduce clumping for increased growth reproducibility (46)) serving as a comparison (**Figure 2A),** to visualize relative differences in metabolic capabilities. From this subset, all grew on methanol, confirming the initial growth phenotype, and all strains also grew on ethanol and methylamine. SLI 231, 233, 499, 505, and 575 grew on vanillic acid. SLI 516, 575, and 576 were the only strains of this subset capable of growth on fructose. Interestingly, these three strains also exhibited biofilm formation during growth in liquid cultures (**Figure 2B, C**), which has been shown in other environmental methylotroph isolates. Biofilm formation in SLI 516, 575, and 576 in various substrates in the presence and absence of LaCl_3_ was verified and quantified via crystal violet assays (data not shown). Cell dry weight measurements of representative SLI strains in all substrates mentioned here in the presence and absence of La^3+^ were determined and found to be similar to cell dry weight of *M. extorquens* AM1 and PA1 (data not shown); thus, growth comparisons across strains are feasible.

### Addition of lanthanides influences the catabolism of alcohols

Based on our initial screen for substrate utilization on solid media, six representative isolates (SLI 231, 233, 384, 499, 505, 575) were chosen as a test cohort for downstream growth analyses because they displayed expanded metabolic capabilities as well as differences in lanthanide-dependent or lanthanide-associated genes, as compared to model organisms *M. extorquens* AM1 or PA1. To investigate the impact of La^3+^ on this collection, the growth rates and final ODs of the test cohort strains were compared to each other and to *M. extorquens* AM1 and PA1 during growth on methanol and ethanol as lanthanides have been shown to influence the oxidation of alcohols in methylotrophic bacteria (18,22).

Growth rates (**Figure 3A**, **3C**) and final ODs (**Figure 3B**, **3D**) on 20 mM methanol and 34 mM ethanol, both with and without La^3+^ in the growth medium, were measured. Growth rates on methanol were significantly higher in the presence of La^3+^ for 4 out of 6 SLI strains (SLI 231, SLI 233, SLI 499, SLI 505; **Figure 3A**) and for *M. extorquens* PA1. In the presence of La^3+^, all SLI strains except SLI 384 had significantly higher growth rates than *M. extorquens* AM1 and were comparable to that of *M. extorquens* PA1 (*p*-values for significance of all strains from one-way ANOVA in **Table S3)**. The addition of lanthanides did not significantly impact the final ODs (**Figure 3B**) obtained for any of the SLI strains. SLI 575 had significantly lower final ODs in the presence of absence of La^3+^ as compared to all other strains, though this could be an artifact of biofilm formation (**Figure 2C**) preventing accurate OD_600_ readings.

**Figure 3.**
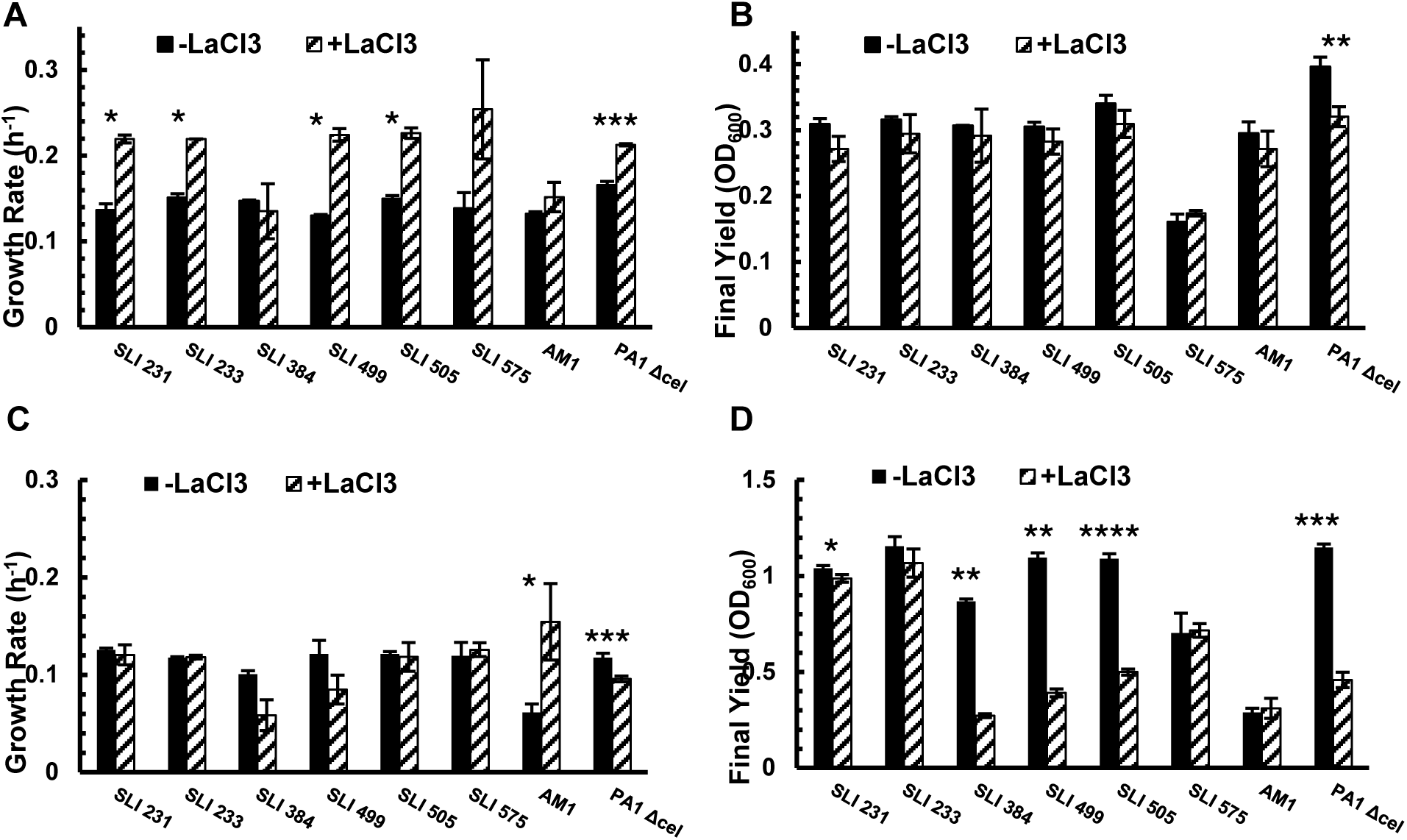
Growth phenotypes of SLI test cohort on alcohols in the presence and absence of La^3+^. **A.** Growth rates on 20 mM methanol +/- 2 µM LaCl_3_. **B.** Final ODs on 20 mM methanol +/- 2 µM LaCl_3_. **C.** Growth rates on 34 mM ethanol +/-2 µM LaCl_3_. **D.** Final ODs on 34 mM ethanol +/- 2 μM LaCl_3_. Black bars (left) represent cultures grown in the absence of LaCl_3_; hatched bars (right) represent cultures grown in the presence of LaCl_3_. N=2-4. Error bars represent standard deviation. Significant differences for each strain +/- LaCl_3_ determined by Student’s paired t-test (*, p<0.05; **, p<0.01; ***, p<0.001; ****, p<0.0001). Significant differences between all strains +/- LaCl_3_ determined using one-way ANOVA with post-hoc Tukey HSD (**Table S3)**

Growth rates on ethanol (**Figure 3C**) were not impacted by the presence of La^3+^ for any of the SLI strains, but were significantly higher for *M. extorquens* AM1 and significantly lower for *M. extorquens* PA1; the physiology behind the opposite phenotypes in these two strains is unknown. In the absence of La^3+^, all SLI strains and *M. extorquens* PA1 had significantly higher growth rates than *M. extorquens* AM1. Strains encoding the same set of lanthanide-dependent alcohol dehydrogenases exhibited drastically different final ODs on ethanol in the presence and absence of La^3+^ (**Figure 3D**). SLI 384, 499, 505, and *M. extorquens* PA1 had more than double the final OD in the absence of La^3+^. *M. extorquens* AM1 had low final ODs regardless of La^3+^. SLI 231, 233, and 575 were similar to *M. extorquens* AM1 in that the presence of La^3+^ did not affect their final OD, yet all strains had higher final ODs than *M. extorquens* AM1.

Despite SLI 231, 233, 499, 505, and *M. extorquens* AM1 all encoding the same lanthanide-dependent alcohol dehydrogenases (XoxF1, XoxF2, ExaF) and having greater than 97% genomic similarity, these strains exhibit growth phenotypes on alcohols in the presence or absence that are significantly different from each other. More SLI strains phenotypically mimic *M. extorquens* PA1 than *M. extorquens* AM1 despite most SLI strains of the test cohort encoding lanthanide-associated genes similar to *M. extorquens* AM1’s repertoire. Thus, the extent to which lanthanide-dependent enzymes affect alcohol utilization remains unknown, and genetically similar strains do not have predictable phenotypes based on genomic analysis or comparison to model organisms alone.

### Robust aromatic acid utilization by SLI strains reveals novel lanthanide-related phenotypes

Recently, facultative methylotrophy has expanded to include species capable of growth on methoxylated aromatic acids within the *nodulans* and *aquaticums* clades of *Methylobacterium* (12). Genes that confer the ability to grow on methoxylated aromatic acids (ferulic acid, vanillic acid, protocatechuic acid, *p-*hydroxybenzoic acid) exist as a horizontally-transferred genetic island that is notably absent from the *extorquens* clade (12), save for their presence in *Methylobacterium sp.* AMS5 which was isolated from a soybean stem and shares many similarities to *M. extorquens* AM1 and PA1 (47) (see **Table 1** for genomic comparison). Methylotrophic growth on methoxylated aromatic acids is predicted to occur via an initial demethoxylation of the methoxy group that is released as formaldehyde to generate protocatechuic acid; formaldehyde can be assimilated or dissimilated via methylotrophic pathways and protocatechuic acid can be converted to acetyl-CoA and succinyl-CoA for assimilation via common heterotrophic pathways (12). Our screen isolated 78 strains in the *extorquens* clade capable of growth on the methoxylated aromatic acid, vanillic acid. Of the SLI strains with assembled genomes, those capable of growth on vanillic acid encode gene islands identical to each other (**Table S2**). Aromatic acid gene islands in SLI strains include all of the genes found in the aromatic acid gene island in *Methylobacterium sp.* AMS5 although specific regulatory genes are in a different order. To demonstrate the robustness of SLI growth on aromatic acids, SLI strains from the test cohort were grown on a low (5 mM) and high (12 mM) concentration of vanillic acid (22,29,30) in the presence and absence of LaCl_3_ and their growth was compared to that of *Methylobacterium sp.* AMS5 (**Figure 4**). SLI 384 from the test cohort as well as *M. extorquens* AM1 and *M. extorquens* PA1 Δ*cel* were used as controls, as these strains do not encode genes for vanillic acid catabolism and therefore do not grow on vanillic acid (**Figure 2**).

**Figure 4.**
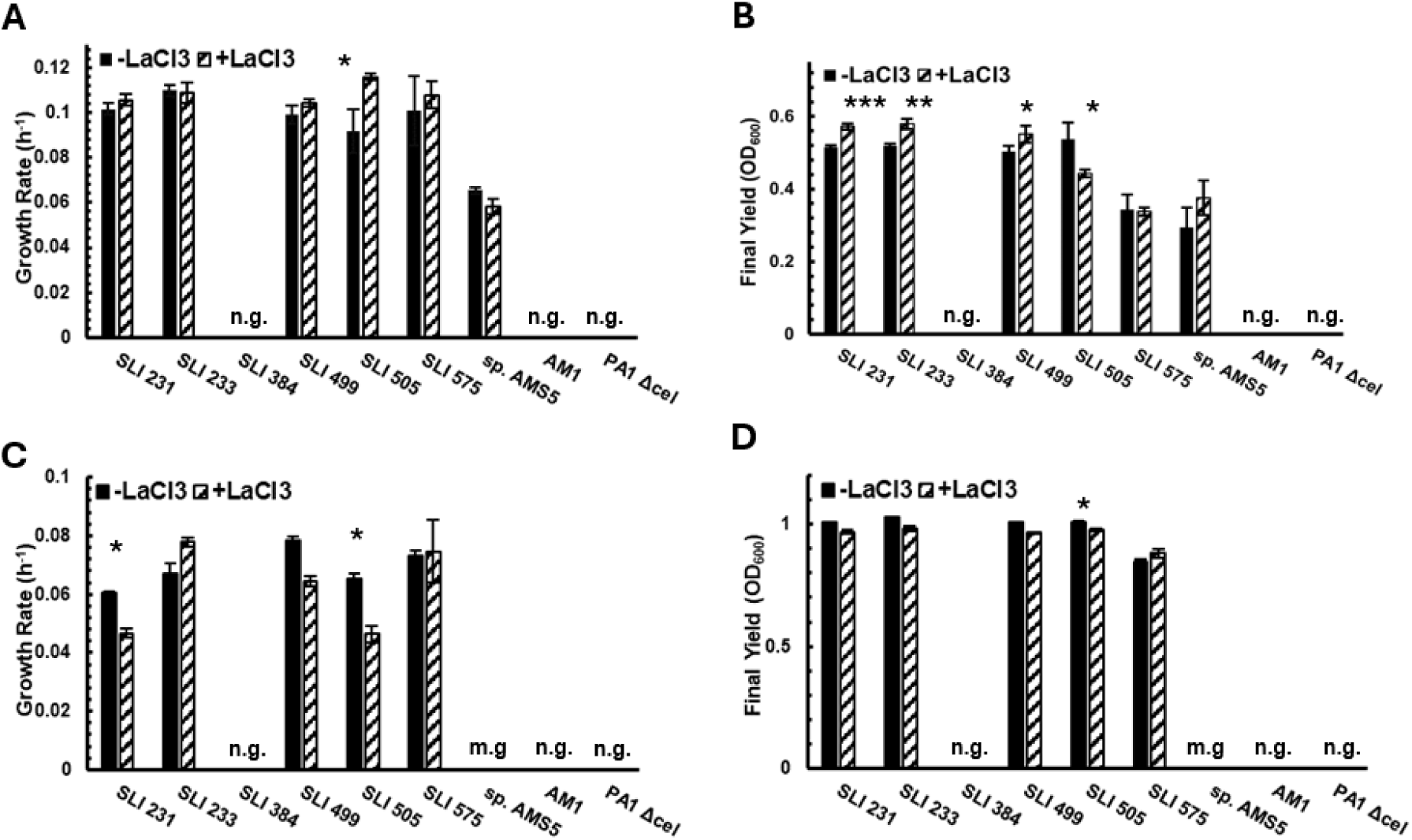
Growth phenotypes of SLI test cohort, *Methylobacterium sp.* AMS5, *M. extorquens* AM1, and *M. extorquens* PA1 Δ*cel* on vanillic acid in the presence and absence of La^3+^. **A** Growth rates on 5 mM vanillic acid +/- 2 μM LaCl_3_. **B.** Final ODs on 5 mM vanillic acid +/- 2 μM LaCl_3_. **C.** Growth rates on 12 mM vanillic acid +/- 2 μM LaCl_3._ **D.** Final ODs on 12 mM vanillic acid +/- 2 μM LaCl_3_. n.g., no growth; m.g., minimal growth (<OD_600_ 0.2). Black bars (left) represent cultures grown in the absence of LaCl_3_; hatched bars (right) represent cultures grown in the presence of LaCl_3_. N=2-4. Error bars represent standard deviation. Significant differences for each strain +/- LaCl_3_ determined by Student’s paired t-test (*, p<0.05; **, p<0.01; ***, p<0.001). Significant differences between all strains +/- LaCl_3_ determined using one-way ANOVA with post-hoc Tukey HSD (**Table S3**)

Two striking phenotypes emerge when comparing growth on different concentrations of vanillic acid. First, all strains exhibit slower growth rates on high concentrations of vanillic acid compared to low concentrations of vanillic acid (**Figure 4A, C**). Despite encoding nearly identical aromatic acid gene islands, *Methylobacterium sp.* AMS5 exhibits poor growth on high concentrations of vanillic acid compared to the SLI strains (**Figure 4C)** and there are significant differences in growth between the SLI strains as well (*p*-values for significance of all strains from one-way ANOVA in **Table S3**). Second, addition of lanthanides influences aromatic acid utilization based on substrate concentration despite all SLI strains encoding identical aromatic acid gene islands. SLI 505 has higher growth rates with lanthanides on 5 mM vanillic acid but lower growth rates with lanthanides on 12 mM vanillic acid (**Figure 4A, C)**. SLI 231 has lower growth rates with lanthanides only during growth on 12 mM vanillic acid, a trend also seen in SLI 499 although the differences are not significant (**Figure 4C**). Conversely, SLI 233 has marginally higher growth rates during growth on 12 mM vanillic acid **(Figure 4C**). Lanthanides variably affect final ODs during growth on low concentrations of vanillic acid (**Figure 4B**), but have little effect on final ODs during growth on high concentrations of vanillic acid (**Figure 4D**).

### SLI strains encode identical methylamine oxidation genes but display different growth phenotypes

Lanthanide utilization during alcohol oxidation in methylotrophs has been well-characterized (18,34,37), yet methylotrophs can also take advantage of other substrates in the phyllosphere whose metabolism may or may not be influenced by lanthanides. For example, methylotrophs have several mechanisms by which they can grow on the C_1_ compound, methylamine. Methylamine dehydrogenases (*mauFBEDACJGLMN*) catalyze the oxidation of methylamine to formaldehyde, which can be further oxidized to formate for assimilation or dissimilation (8,42,48). Alternatively, methylamine can be converted via the *N*-methylglutamate pathway (*mgdDCBA*, *mgsABC*, *gmaS*) to methylene-tetrahydrofolate for assimilation or dissimilation, although the extent of formaldehyde as an obligate intermediate in this pathway remains unknown (42). The model organism *M. extorquens* AM1 encodes both pathways yet primarily uses methylamine dehydrogenase during growth on methylamine, whereas the model organisms *M. extorquens* PA1 and *Methylobacterium sp.* AMS5 only encode the N-methylglutamate pathways (48). Organisms solely utilizing the *N*-methylglutamate pathway have markedly slower growth on methylamine than those that can catalyze the direct oxidation to formaldehyde (46,48). All SLI genomes encode only the genes for the *N*-methylglutamate pathway for methylamine utilization similar to what is found in *M. extorquens* PA1.

To compare the growth of SLI strains on methylamine to *M. extorquens* AM1 and *M. extorquens* PA1 Δ*cel*, all strains were grown in 15 mM methylamine in the presence and absence of LaCl_3_. Growth rates (**Figure 5A**) and final ODs (**Figure 5B**) are reported. Addition of LaCl_3_ did not significantly impact the growth rate or yield for any SLI strain tested. Yet, *M. extorquens* PA1 had significantly higher growth rates and lower final ODs in the presence of lanthanides; this is the first report of lanthanides influencing methylamine metabolism, although the difference is subtle. Also of note, SLI 384 had nearly triple the lag time and half the growth rate and final yield as other SLI strains despite encoding identical *N*-methylglutamate pathway genes. This suggests additional details about methylamine catabolism in SLI strains beyond the *N*-methylglutamate pathway that result in the diminished growth phenotypes observed.

**Figure 5.**
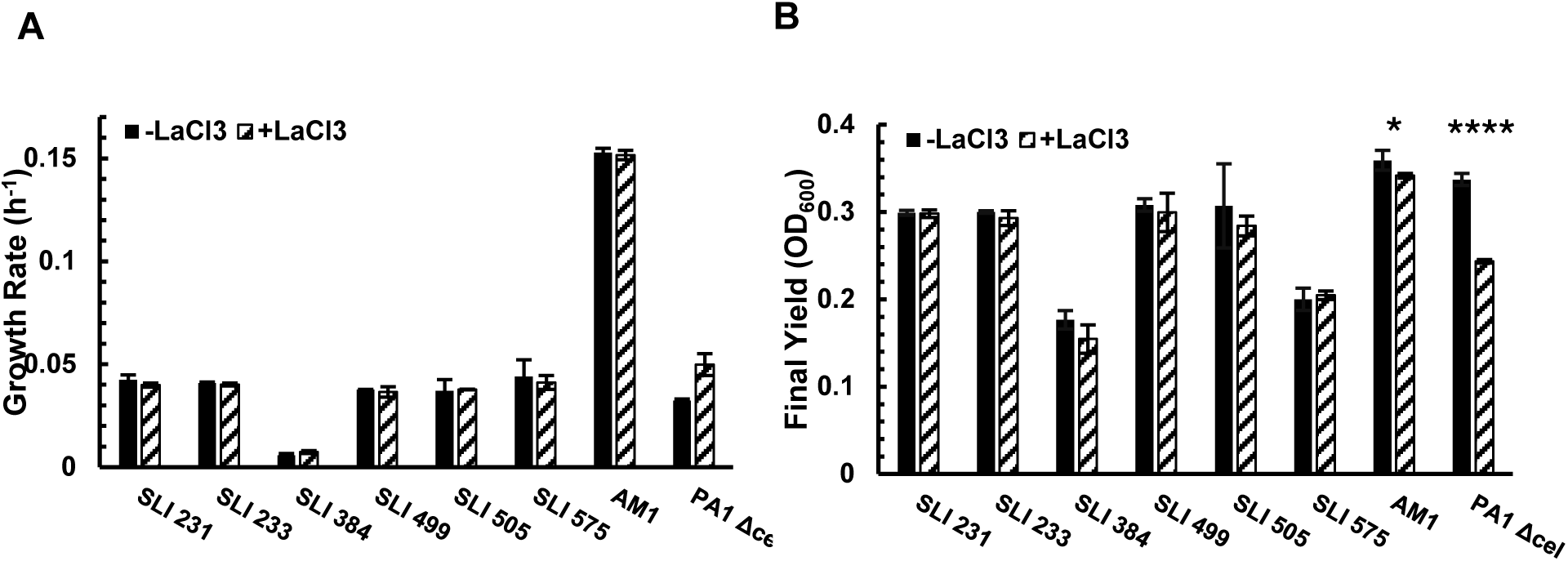
Growth phenotypes of SLI test cohort, *M. extorquens* AM1, and *M. extorquens* PA1Δ*cel* on methylamine in the presence and absence of La^3+^. **A.** Growth rates on 15 mM methylamine +/- 2 μM LaCl_3_. **B.** Final ODs on 15 mM methylamine +/- 2 μM LaCl_3_. Black bars (left) represent cultures grown in the absence of LaCl_3_; hatched bars (right) represent cultures grown in the presence of LaCl_3_. N=2-4. Error bars represent standard deviation. Significant differences for each strain +/- LaCl_3_ determined by Student’s paired t-test (*, p<0.05; **, p<0.01; ***, p<0.001; ****, p<0.0001). Significant differences between all strains +/- LaCl_3_ determined using one-way ANOVA with post-hoc Tukey HSD (**Table S3**)

### SLI strains expand fructose utilization in the extorquens clade

Most methylotrophs, including *M. extorquens* AM1 and PA1, encode all of the genes necessary for sugar oxidation but lack sugar-specific transporters and assimilatory pathways required for their utilization as a primary substrate (49). Notably, complete sugar catabolism has only been demonstrated in endophytic *extorquens* clade members (50). Thus, it was striking to isolate ten strains capable of robust growth on fructose as the sole carbon source (**Table 2**). Of the strains with assembled genomes, SLI 516, SLI 575, and SLI 576 were capable of growth on fructose. Interestingly, these three SLI strains also displayed the lowest sequence similarity to other SLI strains and to *M. extorquens* AM1 and PA1 based on ANI (**Figure 1B**), hinting that the differences in genomes might be due in part to unknown genes specific to sugar transport and catabolism. To quantify growth on fructose, SLI 575 was grown on 25 mM fructose in the presence and absence of LaCl_3_ (**Figure 6**). Growth rates and final ODs in both conditions were nearly identical. Growth rates on fructose were comparable to rates for methylamine and substantially lower than rates for methanol or vanillic acid. To our knowledge, SLI 516, 575, and 576 are the first identified epiphytic *extorquens* strains capable of growth on a sugar substrate.

**Figure 6.**
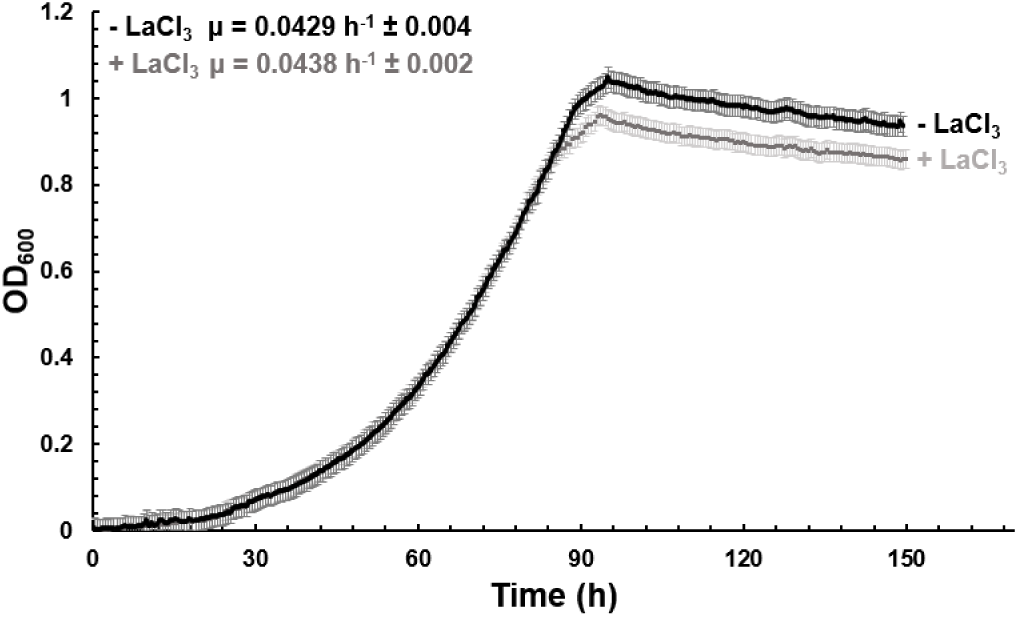
Growth curves of SLI 575 on 25 mM fructose +/- 2 μM LaCl_3_. N=4. Error bars represent standard deviation

## Discussion

Here, we describe an enrichment of *Methylobacterium* isolated in a lanthanide-dependent manner to characterize the influence of lanthanides during the metabolisms of natural methylotrophic populations in soybean plants, revealing novel insights into methylotrophic biodiversity, metabolic capabilities, and lanthanide-dependent metabolisms. Methylotrophs that inhabit the phyllosphere influence both the plants that they colonize and global biogeochemical processes (1,2,2,7,51). Yet, studies investigating the physiology, biodiversity, and ecology of methylotrophic communities of the phyllosphere have only emerged within the last decade (6,13,52) and rarely take into account the role of lanthanides in influencing their community composition. Furthermore, despite the abundance of methylotrophs in the phyllosphere (6,7,53), the bulk of phylogenomic studies on *Methylobacterium* species use genomes isolated from non-phyllosphere environments.

Marker gene sequencing for assigning taxonomic groups has proven difficult in *Methylobacterium*, as16S rRNA has poor phylogenetic resolution in *Methylobacterium* (38). We characterized our soybean methylotroph enrichment collection using *rpoB* (**Figure 1A**), a highly polymorphic single-copy gene with sub-species resolution within *Methylobacterium*(38). Our phylogenetic analysis revealed two insights: 1) all of our isolates are closely related to each other (**Figure 1A**) and fall within the *extorquens* clade of *Methylobacterium* (**Figure 1A, B**), and 2) even amongst nearly identical strains (**Figure 1B**), there is broad diversity in terms of the possession of lanthanide-related genes and metabolic capabilities, and that these differences are often reflected in how strains group based on *rpoB* sequences. Overall, our results substantiate *rpoB* as a superior marker gene for phylogenetic analysis of *Methylobacterium* species, and, importantly, highlight how phenotypic characterizations can reveal novel strains that appear identical by *rpoB* sequence alone.

Crops, such as soybeans, can sequester lanthanides from the soil through repeated cycles of harvest (Colleen Doherty, unpublished data/ personal communication), limiting lanthanide availability in the soil and affecting lanthanide availability for methylotrophs in the phyllosphere. Thus, it is not surprising that our microbial isolation from soybean plants did not reveal any novel lanthanide-dependent phenotypes despite previous studies identifying organisms that have increased heterogeneity in the presence of lanthanides (54) or that are strictly dependent on lanthanides for growth (34,36,37,55,56). Our isolation media lacked PQQ, constraining the isolation to methylotrophs capable of synthesizing this cofactor for use in alcohol dehydrogenases (15). Future isolations using media supplemented with this cofactor might yield greater diversity of isolates. However, cultivation-independent proteogenomic characterizations of the soybean phyllosphere identified high degrees of similarity in *Methylobacterium* populations on the plant as compared to reference *Methylobacterium* strains (6,53), further underlying the abundance of *Methylobacterium* species in the phyllosphere environment. The low species diversity of our enrichment collection provides a model by which to interrogate how closely related strains restrict resource overlap to co-exist in soybean leaf environments.

The maintenance of both calcium-dependent and lanthanide-dependent methanol dehydrogenase systems in our SLI strains could suggest that 1) lanthanides are not always present or bioavailable in the phyllosphere and therefore MxaFI remains essential for methylotrophic growth; 2) lanthanides are present but extensively competed for, so calcium-dependent methanol oxidation systems offer an alternative for methylotrophic growth; and/or 3) lanthanide-dependent enzymes have yet-uncharacterized physiological or regulatory roles that necessitate their maintenance. These hypotheses could be examined further by repeating our isolations in non-domesticated phyllosphere environments where lanthanides are more prevalent or bioavailable and where the isolation of obligately lanthanide-dependent methylotrophs might be more likely.

Our isolates include strains that encode all or a subset of known lanthanide-dependent (XoxF1, XoxF2, ExaF) and lanthanide-binding (lanmodulin) proteins (**Figure 1** and **Table S2**), yet display different phenotypes during lanthanide-dependent growth on ethanol (**Figure 3**) that cannot be correlated to the presence or absence of these genes alone. A previous study comparing *M. extorquens* AM1 and *M. extorquens* PA1 on 7.5 mM ethanol in the absence of lanthanides (46) found that *M. extorquens* AM1 had poorer growth than PA1, similar to the trends observed in our study. The authors hypothesized that this difference might be due to an aldehyde dehydrogenase (Mext_1295) present in PA1 but absent in AM1. As homologs of this gene were found to be absent in all of our SLI strains, further investigations into the optimal substrate concentration range for ethanol growth of each SLI strain, as well as comparisons of promoter and/or regulatory regions related to ethanol metabolism genes are necessary to understand this striking ethanol phenotype.

SLI strains lacking XoxF2, ExaF, and lanmodulin emerge as an interesting model by which to interrogate the role of various lanthanide-related proteins in methylotrophs both in laboratory and plant environments, as compared to model organisms like *M. extorquens* AM1 or PA1. Lanmodulin is reported to be one of the most highly expressed peptides in the phyllosphere (29), has been proposed to function as a lanthanide biosensor, and has been shown to bind lanthanides(45), yet its functional role in lanthanide metabolism remains elusive. A previous study in *Methylobacterium aquaticum* 22A (57), which natively encodes *lanM*, found that overexpression of *lanM* results in faster growth on methanol with 20 nM LaCl_3_ and slower growth with 100 nM LaCl_3_ (33). Future work expressing *lanM* in SLI strains lacking lanmodulin during growth on methanol and ethanol with different concentrations and species (soluble vs poorly soluble) of lanthanides and accompanying metal analysis via ICP-MS could be revealing for identifying putative roles for lanmodulin in the phyllosphere.

Previous studies have investigated methylamine metabolism in the model organisms *M. extorquens* AM1 and PA1 (42,48), yet the influence of lanthanides on this metabolism has yet to be reported. Here, we demonstrate that our SLI strain exhibit the same phenotypes in methylamine regardless of LaCl_3_ addition (**Figure 5**), in contrast to novel lanthanide-dependent phenotypes observed in *M. extorquens* AM1 and PA1, but the exact role of lanthanides in this metabolism remain unknown. Interestingly, SLI 384 and SLI 223 (data not shown for the latter) have much slower growth in methylamine than all other SLI strains; this suggests metabolic bottlenecks towards efficient methylamine assimilation that are not immediately obvious from a preliminary genomic analysis of C_1_ assimilation genes. One possibility is that methylamine that is incorporated via the N-methylglutamate pathway preferentially serves as a nitrogen source rather than a carbon source (42) but future studies are required to validate this hypothesis in our SLI strains.

Although the ability to utilize methanol as a substrate has provided a competitive advantage for methylotrophs in the phyllosphere (6,7,58), sugars and aromatic acid are abundant enough to support the growth of diverse microorganisms (3,12) It is therefore advantageous for microbes to be able to utilize both multicarbon sources, such as fructose and vanillic acid, and single carbon sources like methanol (5,59). Detailed investigations into methylotrophic aromatic acid catabolism in the presence and absence of lanthanides is necessary to understand how this metabolism functions in carbon cycling of lignin by-products in natural environments, as well as identify additional roles for lanthanides in the metabolism of non-alcohol substrates. The decrease in growth rates in the presence of lanthanides only during growth on high concentrations of aromatic acid is surprising (**Figure 4**), considering that all known lanthanide-dependent enzymes are in the periplasm yet all enzymes used for aromatic acid utilization are in the cytoplasm; the role of lanthanides during this metabolism is an ongoing area of research. The substantial final OD (**Figure 4B**, **4D)** but decreased growth rate **(Figure 4A**, **4C)** achieved by SLI strains on high concentrations vs low concentrations of vanillic acid demonstrates that the ability to grow on methoxylated aromatic acids is widely distributed among members of the *extorquens* clade, more robust than what has been reported in other model strains, and could reveal novel insights about lanthanide metabolism.

Genomic analyses for fructose transporters and assimilatory genes in SLI 516, 575, or 576 (SLI strains with assembled genomes capable of growth on fructose) identified a gene cluster annotated with genes for fructose-specific transporters homologous to HPr and Enzyme II of the canonical phosphotransfer system in *Escherichia coli*(60,61). Genes homologous to Enzyme I were not found in this cluster, but SLI 516, 575, and 576 all encoded genes for phosphofructokinase-1, which regulates glycolysis and converts fructose-6-phosphate to fructose 1,6-bisphosphate(61), and a porin in the same cluster (**Table S2**). For comparison, genes homologous to HPr, Enzyme I, and Enzyme II from *E. coli* were also found in *M. extorquens* AM1 and PA1 and all other SLI strains, but they were not sugar-specific and growth on glucose or fructose is not possible in these strains.

Taken together, the methylotrophic community in the phyllosphere of soybeans is metabolically diverse and heavily influenced by lanthanide availability when consuming methanol. We hypothesize that the capacity to expand substrate repertoires might also be important drivers of efficient colonization(5,58,59). Whether the selection for low species diversity is at the level of the plant or arises through natural selection within methylotrophic communities, this finding points to the evolution of niche partitioning strategies among a single species in order to maximize resources in shared habitats (5,52,62). Results from this study pave the way for future work to investigate (1) expanded metabolic capabilities in *extorquens* clade methylotrophs, (2) tradeoffs associated with lanthanide-dependent and -independent alcohol metabolism, and (3) niche partitioning among genetically identical strains during plant colonization.

## Materials & Methods

### Methylotroph isolation

Environmental methylotrophic bacterial strains were isolated from soybean plants (*Glycine max*) growing on the Michigan State University Agronomy Farm (East Lansing, Michigan, U.S.A. 42.6908, -84.4866). Four leaves were harvested from four different plants on September 7, 2018, and each leaf was placed into a sterile 50 mL conical tube. 50 ml of sterile 50 mM phosphate buffer (pH 7.3) was added to each tube, the tubes were vortexed for 5 minutes, and 100 μL of buffer was spread onto solidified (1.5% wt/vol agar) MP minimal salts media (33) <u>optimized for robust and reproducible growth of *Methylobacterium* strains</u>. The isolation medium contained 125 mM methanol, RPMI 1640 Vitamins Solution (Sigma Aldrich, St. Louis, MO, USA) diluted to 1X, and 50 μg/mL cycloheximide. The vitamins solution contains all twenty amino acids, D-biotin, choline chloride, folic acid, myo-inositol, niacinamide, p-amino benzoic acid, D-pantothenic acid, pyridoxal HCl, pyridoxine HCl, thiamine HCl, vitamin B12, KH_2_PO_4_, NaCl, and Na_2_HPO_4_ (exact composition on manufacturer’s website) and has been shown to enhance growth of environmental methylotroph isolates. The cycloheximide prevents isolation of methylotrophic fungi which are outside of the scope of this study. Two sets of isolation media were prepared; one with and one without 2 µM LaCl_3_. Isolation plates were grown at 30 °C for several days until bacterial colonies were visible. Pink colonies were selected from the plates and streaked for isolated colonies onto solidified media of the same composition to confirm growth. Single colonies of isolated strains were then inoculated into 650 μL of MP medium with 125 mM methanol and 1X RPMI 1640 Vitamins Solution in a 48 well microplate. All isolated strains were inoculated into medium with the same concentration of La^3+^ as they were isolated with, and then incubated shaking for 48 hours at 200 rpm on an Innova 2300 platform shaker (Eppendorf, Hamburg, Germany) at 30°C. Strains were frozen for future use by adding 25 μL of sterile dimethyl sulfoxide to 500 μL of late-exponential phase culture in a sterile screw-cap vial, flash-freezing in liquid nitrogen, and storing at -80°C.

### Strains cultivation

Cultures used for growing and maintaining strains, extracting DNA, and subculturing for growth phenotypic analyses were prepared as follows: Freezer stocks of each strain were streaked out onto MP agar supplemented with 15 mM succinate and 1X RPMI 1640 Vitamins Solution and incubated at 30°C until single colonies emerged (2-4 days). Single colonies of each strain were used to inoculate 3 mL cultures of MP with 15 mM succinate and 1X RPMI 1640 Vitamins Solution and grown overnight in round-bottom plastic culture tubes (ThermoFisher Scientific, Waltham, MA, USA) at 30°C, shaking at 200 rpm on an Innova S44i shaker (Eppendorf, Hamburg, Germany) to an OD_600_ of approximately 1.5. All media was prepared in glassware that was biologically cleaned with an obligately lanthanide-dependent strain, to ensure that glassware is lanthanide-free. The three transfers in lanthanide-free medium from freezer stocks to streaking for single colonies to overnight culture outgrowth in lanthanide-free media and glassware have been shown to be sufficient for eliminating lanthanide carryover and cell-sequestration (63). See sections below for downstream analyses performed on these cultures.

### Chromosomal DNA extraction for whole-genome and marker gene sequencing

Genomic DNA was extracted from 3 mL overnight cultures of each strain (see ***Strains Cultivation***) using the DNeasy PowerSoil Pro Kit from Qiagen (Hilden, Germany) per manufacturer’s protocol. Genomic DNA was quantified using a Take3 Microvolume Plate Spectrophotometer (BioTek, Winooski, VT, USA). Whole-genome sequencing via PacBio Sequel II was performed by the Department of Energy’s Joint Genome Institute (JGI, Walnut Creek, CA, USA). Genomes can be accessed through JGI Integrated Microbial Genomes & Microbiomes (IMG) (see **Table 1** for IMG Genome IDs). Marker gene sequencing was performed by PCR amplifying regions of *16s* rRNA or *rpoB* using universal primers^18^ (16s_fwd: 5’GAGTTTGATCCTGGCTCA3’, 16s_rev: 5’TACCTTGTTACGACTT3’; rpoB_fwd: 5’AAGGACATCAAGGAGCAGGA3’, rpoB_rev: 5’ACSCGGTAKATGTCGAACAG3’), confirming products via gel electrophoresis on a 1% agarose gel, purifying PCR products using the GeneJET PCR Purification Kit (ThermoFisher Scientific, Waltham, MA, USA), and Sanger sequencing the PCR products. Untrimmed sequences were aligned using MAFFT and MUSCLE, and a Maximum-Likelihood tree using a Tamura-Nei model and 100 bootstrap replications was constructed using MEGA. Average nucleotide identity scores were calculated through JGI IMG interface for pairwise genome comparisons using whole-genome sequences from the IMG database as inputs (see **Table 1** for JGI IMG Genome IDs).

### Growth phenotypic analyses

Isolate strains were cultivated as described in ***Strains Cultivation*** and were screened for their ability to grow on diverse substrates in the presence and absence of La^3+^. To screen for the ability to grow on diverse substrates, overnight cultures of each strain were washed twice in MP media with no carbon source at 2000 x g for 10 minutes, resuspended in MP media to an OD600 of 0.1, and spotted (10 uL) onto MP agar plates supplemented with 1X 1640 RPMI Vitamins Solution, and either a soluble substrate (20 mM methanol, 34 mM ethanol, 15 mM methylamine, 5 mM potassium oxalate, 25 mM glucose, 25 mM fructose) or an insoluble substrate (5 mM vanillic acid); the insoluble substrates were added as non-sterilized powders directly to the melted agar prior to pouring the plate. Agar plates were incubated at 30°C for 2-4 days or until visible colonies formed.

To screen for La^3+^ effects during growth on methanol, overnight cultures of each strain were washed twice in MP media with no carbon source at 2000 x g for 10 minutes and inoculated at an OD600 of 0.1 into a transparent 96-well plate (Corning, Corning, NY, USA) containing 200 uL of MP media supplemented with 125 mM methanol, 1X RPMI 1640 Vitamins Solution, and +/- 2 μM LaCl_3_ for endpoint growth assays. Strains with demonstrated growth on solid media of different substrates were additionally phenotyped in liquid media in the presence and absence of La^3+^. Overnight cultures of strains of interest were washed twice in MP media with no carbon source at 2000 x g for 10 minutes and inoculated at an OD600 of 0.1 into a transparent 48-well plate (Corning, Corning, NY, USA) containing 650 uL of MP media supplemented with 1X 1640 RPMI Vitamins Solution, +/- 2 μM LaCl_3_, and appropriate carbon source (20 mM methanol, 15 mM methylamine, 34 mM ethanol, 12 mM vanillic acid, 25 mM fructose). All endpoint assays and growth curves were carried out at 30°C and 548 rpm using a Synergy HTX plate reader (BioTek, Winooski, VT, USA) with OD600 readings measured every 30 minutes. Data was analyzed using Microsoft Excel. 2-4 replicates were run for each strain in each condition, and at least 10 exponential-phase data points were used for linear regression analysis to calculate the growth rates of each strain. Paired t-tests were performed to identify statistically significant (p < 0.05) differences between growth in the presence or absence of LaCl_3_ for each strain. One-way ANOVA with post-hoc Tukey HSD was performed to identify statistically significant (p < 0.05) differences among all strains in the presence or absence of LaCl_3._.

## Acknowledgements

We thank Allison Hunt and Zachary Jensen—previous members of the Martinez-Gomez Lab— for their invaluable efforts during the early stages of this project. We would also like to acknowledge Jean-Baptiste Leducq, Christopher Marx, Alexander Alleman, and all members of the Marx Lab at the University of Idaho for their thoughtful comments and suggestions during experimental planning and manuscript drafting. The information, data, or work presented herein was funded by the United States Department of Energy, Office of Science, Office of Biological and Environmental Research, under Award Number DE-SC0022318 subaward SH5849-772894; by the National Science Foundation under Grant 2142154; by the National Science Foundation under Grant 2127732. A.M.G. was supported in part by the National Institutes of Health Genetic Dissection of Cells and Organisms Training Grant 1T32GM132022-01.

## Data Availability

The data generated and/or analyzed during the current study as well as stocks of novel reported strains are available from the corresponding author upon reasonable request. Genomes from novel strains reported in this study can be found through JGI IMG using the IMG Genome IDs indicated in **Table 1**.

