## Supplemental Table 1 for "Lanthanide-dependent isolation of phyllosphere methylotrophs selects for a phylogenetically conserved but metabolically diverse community"

**Table S1.** Average nucleotide identity of all 11 SLI strains compared to evolutionarily-related methylotroph and non-methylotrophic species, including all species represented in phylogenetic tree from **Figure 1.** A, *Afipia felis* 76713; B, *Bradyrhizobium diazoefficiens* USDA 110; C, *Methylobacterium aquaticum* DSM 16371; D, *Methylobacterium aquaticum* MA-22A; E, *Methylobacterium brachiatum* 111MFTsu3.1M4; F, *Methylobacterium chloromethanicum* CM4; G, *Methylobacterium extorquens* AM1; H, *Methylobacterium extorquens* DM4; I, *Methylobacterium extorquens* PA1; J, *Methylobacterium indicum* NS230; K, *Methylobacterium indicum* SE2,11; L, *Methylobacterium komagatae* DSM 19563; M, *Methylobacterium nodulans* ORS 2060; N, *Methylobacterium organophilum* DSM 760; O, *Methylobacterium oryzae* CBMB20; P, *Methylobacterium phyllosphaerae* CBMB27; Q, *Methylobacterium platani* JCM 14648; R, *Methylobacterium populi* BJ001; S, *Methylobacterium pseudosasicola* BL36; T, *Methylobacterium radiotolerans* JCM 2831; U, *Methylobacterium* sp. 275MFSha3.1; V, *Methylobacterium* sp. 4-46; W, *Methylobacterium* sp. AMS5; X, *Methylobacterium* sp. UNCCL125; Y, *Methylobacterium* sp. WSM2598; Z, *Methylobacterium* tarhaniae DSM 25844; AA, *Methylobacterium* variabile DSM 16961; AB, *Methylobacterium* zatmanii PSBB041; AC, *Pseudomonas putida* KT2440; AD, *Rhodopseudomonas palustris* DSM 126; AE, SLI 223; AF, SLI 231; AG, SLI 233; AH, SLI 274; AI, SLI 285; AJ, SLI 384; AK, SLI 499; AL, SLI 505; AM, SLI 516; AN, SLI 575; AO, SLI 576

| | A | B | C | D | E | F | G | H | I | J | K | L | M | N | O | P | Q | R | S | T | U | V | W | X | Y | Z | AA | AB | AC | AD | AE | AF | AG | AH | AI | AJ | AK | AL | AM | AN | AO |
|---|---|---|---|---|---|---|---|---|---|---|---|---|---|---|---|---|---|---|---|---|---|---|---|---|---|---|---|---|---|---|---|---|---|---|---|---|---|---|---|---|---|
| **A** | 100 | 78 | 75 | 75 | 75 | 75 | 75 | 75 | 76 | 76 | 75 | 75 | 75 | 75 | 75 | 75 | 78 | 76 | 75 | 75 | 75 | 75 | 76 | 75 | 75 | 76 | 76 | 75 | 72 | 78 | 76 | 76 | 75 | 75 | 75 | 76 | 76 | 76 | 75 | 75 | 75 |
| **B** | 78 | 100 | 76 | 76 | 76 | 76 | 76 | 76 | 76 | 76 | 76 | 75 | 76 | 76 | 75 | 75 | 78 | 76 | 76 | 76 | 76 | 76 | 76 | 75 | 76 | 76 | 76 | 76 | 72 | 80 | 76 | 76 | 76 | 76 | 76 | 76 | 76 | 76 | 76 | 76 | 76 |
| **C** | 75 | 76 | 100 | 89 | 79 | 80 | 79 | 80 | 79 | 89 | 89 | 79 | 82 | 80 | 79 | 79 | 93 | 80 | 79 | 79 | 79 | 82 | 79 | 79 | 82 | 91 | 89 | 79 | 73 | 76 | 79 | 79 | 79 | 79 | 79 | 79 | 79 | 79 | 79 | 79 | 79 |
| **D** | 75 | 76 | 89 | 100 | 79 | 79 | 79 | 80 | 79 | 79 | 93 | 79 | 82 | 80 | 79 | 79 | 93 | 80 | 79 | 80 | 80 | 83 | 79 | 80 | 83 | 90 | 89 | 79 | 73 | 76 | 79 | 79 | 79 | 79 | 79 | 79 | 79 | 79 | 79 | 79 | 79 |
| **E** | 75 | 76 | 79 | 79 | 100 | 80 | 80 | 80 | 80 | 79 | 80 | 81 | 79 | 86 | 86 | 86 | 84 | 80 | 92 | 86 | 86 | 79 | 80 | 86 | 79 | 79 | 80 | 80 | 72 | 76 | 80 | 80 | 80 | 80 | 80 | 80 | 80 | 80 | 80 | 80 | 80 |
| **F** | 75 | 76 | 80 | 79 | 80 | 100 | 97 | 97 | 97 | 79 | 79 | 80 | 79 | 80 | 80 | 80 | 84 | 90 | 80 | 80 | 80 | 79 | 95 | 80 | 79 | 79 | 80 | 97 | 72 | 76 | 97 | 97 | 97 | 97 | 97 | 97 | 97 | 97 | 93 | 93 | 93 |
| **G** | 75 | 76 | 79 | 79 | 80 | 97 | 100 | 97 | 97 | 79 | 79 | 80 | 79 | 80 | 80 | 80 | 84 | 90 | 80 | 80 | 80 | 79 | 95 | 80 | 79 | 79 | 80 | 97 | 72 | 76 | 100 | 97 | 97 | 97 | 97 | 100 | 97 | 97 | 93 | 93 | 93 |
| **H** | 75 | 76 | 80 | 79 | 80 | 97 | 97 | 100 | 97 | 79 | 79 | 80 | 79 | 80 | 80 | 80 | 84 | 90 | 80 | 80 | 80 | 79 | 95 | 80 | 79 | 79 | 80 | 97 | 72 | 76 | 97 | 97 | 97 | 97 | 97 | 97 | 97 | 97 | 93 | 93 | 93 |
| **I** | 76 | 76 | 79 | 79 | 80 | 97 | 97 | 97 | 100 | 79 | 79 | 79 | 79 | 80 | 80 | 80 | 84 | 90 | 80 | 80 | 80 | 79 | 95 | 80 | 79 | 79 | 79 | 97 | 72 | 76 | 97 | 97 | 97 | 97 | 97 | 97 | 97 | 97 | 93 | 93 | 93 |
| **J** | 76 | 76 | 89 | 93 | 79 | 79 | 79 | 79 | 79 | 100 | 99 | 79 | 83 | 80 | 79 | 79 | 93 | 80 | 79 | 80 | 79 | 83 | 79 | 79 | 83 | 90 | 89 | 79 | 73 | 76 | 79 | 79 | 79 | 79 | 79 | 79 | 79 | 79 | 79 | 79 | 79 |
| **K** | 75 | 76 | 89 | 79 | 80 | 79 | 79 | 79 | 79 | 99 | 100 | 79 | 83 | 80 | 79 | 79 | 93 | 80 | 79 | 80 | 80 | 83 | 79 | 79 | 83 | 90 | 89 | 79 | 73 | 76 | 79 | 79 | 79 | 79 | 79 | 79 | 79 | 79 | 79 | 79 | 79 |

| | | | | | | | | | | | | | | | | | | | | | | | | | | | | | | | | | | | | | | | | | |
|---|---|---|---|---|---|---|---|---|---|---|---|---|---|---|---|---|---|---|---|---|---|---|---|---|---|---|---|---|---|---|---|---|---|---|---|---|---|---|---|---|---|
| **L** | 75 | 75 | 79 | 79 | 81 | 80 | 80 | 80 | 79 | 79 | 79 | 100 | 78 | 83 | 81 | 81 | 84 | 80 | 81 | 82 | 82 | 78 | 79 | 82 | 78 | 79 | 79 | 79 | 72 | 75 | 79 | 79 | 79 | 79 | 79 | 79 | 79 | 79 | 79 | 79 | 79 |
| **M** | 75 | 76 | 82 | 82 | 79 | 79 | 79 | 79 | 79 | 83 | 83 | 78 | 100 | 79 | 79 | 79 | 87 | 79 | 79 | 79 | 79 | 86 | 79 | 79 | 86 | 82 | 83 | 79 | 72 | 76 | 79 | 79 | 79 | 79 | 79 | 79 | 79 | 79 | 79 | 79 | 79 |
| **N** | 75 | 76 | 80 | 80 | 86 | 80 | 80 | 80 | 80 | 80 | 80 | 83 | 79 | 100 | 91 | 91 | 84 | 81 | 86 | 99 | 95 | 80 | 80 | 91 | 80 | 80 | 80 | 80 | 73 | 76 | 80 | 80 | 80 | 80 | 80 | 80 | 80 | 80 | 80 | 80 | 80 |
| **O** | 75 | 75 | 79 | 79 | 86 | 80 | 80 | 80 | 80 | 79 | 79 | 81 | 79 | 91 | 100 | 99 | 84 | 80 | 86 | 91 | 91 | 79 | 80 | 99 | 79 | 79 | 80 | 80 | 72 | 76 | 80 | 80 | 80 | 80 | 80 | 80 | 80 | 80 | 80 | 80 | 80 |
| **P** | 75 | 76 | 79 | 79 | 86 | 80 | 80 | 80 | 80 | 79 | 79 | 81 | 79 | 91 | 99 | 100 | 84 | 80 | 86 | 91 | 91 | 79 | 80 | 99 | 79 | 79 | 80 | 80 | 73 | 76 | 80 | 80 | 80 | 80 | 80 | 80 | 80 | 80 | 80 | 80 | 80 |
| **Q** | 78 | 78 | 93 | 93 | 84 | 84 | 84 | 84 | 84 | 93 | 93 | 83 | 87 | 84 | 84 | 84 | 100 | 84 | 84 | 85 | 84 | 87 | 84 | 84 | 87 | 93 | 92 | 84 | 75 | 78 | 84 | 84 | 84 | 84 | 84 | 84 | 84 | 84 | 84 | 84 | 84 |
| **R** | 76 | 76 | 80 | 80 | 80 | 90 | 90 | 90 | 90 | 80 | 80 | 80 | 79 | 81 | 80 | 80 | 84 | 100 | 80 | 81 | 80 | 79 | 90 | 80 | 79 | 80 | 80 | 90 | 72 | 76 | 90 | 90 | 90 | 90 | 90 | 90 | 90 | 90 | 90 | 90 | 90 |
| **S** | 75 | 76 | 79 | 79 | 92 | 80 | 80 | 80 | 80 | 79 | 79 | 81 | 79 | 86 | 86 | 86 | 84 | 80 | 100 | 86 | 86 | 79 | 80 | 86 | 79 | 79 | 79 | 80 | 73 | 76 | 80 | 80 | 80 | 80 | 80 | 80 | 80 | 80 | 80 | 80 | 80 |
| **T** | 75 | 76 | 79 | 80 | 86 | 80 | 80 | 80 | 80 | 80 | 80 | 82 | 79 | 99 | 91 | 91 | 85 | 81 | 86 | 100 | 95 | 80 | 80 | 91 | 80 | 80 | 80 | 80 | 73 | 76 | 80 | 80 | 80 | 80 | 80 | 80 | 80 | 80 | 80 | 80 | 80 |
| **U** | 75 | 76 | 79 | 80 | 86 | 80 | 80 | 80 | 80 | 80 | 80 | 82 | 79 | 95 | 91 | 91 | 84 | 80 | 86 | 95 | 100 | 80 | 80 | 91 | 80 | 80 | 80 | 80 | 73 | 76 | 80 | 80 | 80 | 80 | 80 | 80 | 80 | 80 | 80 | 80 | 80 |
| **V** | 75 | 76 | 82 | 83 | 79 | 79 | 79 | 79 | 79 | 83 | 83 | 78 | 86 | 80 | 79 | 79 | 87 | 79 | 79 | 80 | 80 | 100 | 79 | 79 | 79 | 83 | 83 | 79 | 72 | 76 | 79 | 79 | 79 | 79 | 79 | 79 | 79 | 79 | 79 | 79 | 79 |
| **W** | 75 | 76 | 79 | 79 | 80 | 95 | 95 | 95 | 95 | 79 | 79 | 79 | 79 | 80 | 80 | 80 | 84 | 90 | 80 | 80 | 80 | 79 | 100 | 80 | 79 | 79 | 79 | 95 | 72 | 76 | 95 | 95 | 95 | 95 | 95 | 95 | 95 | 95 | 93 | 93 | 93 |

| | | | | | | | | | | | | | | | | | | | | | | | | | | | | | | | | | | | | | | | | | |
|---|---|---|---|---|---|---|---|---|---|---|---|---|---|---|---|---|---|---|---|---|---|---|---|---|---|---|---|---|---|---|---|---|---|---|---|---|---|---|---|---|---|
| **X** | 75 | 76 | 79 | 80 | 86 | 80 | 80 | 80 | 80 | 79 | 79 | 82 | 79 | 91 | 99 | 99 | 84 | 80 | 86 | 91 | 91 | 79 | 80 | 100 | 79 | 79 | 80 | 80 | 73 | 76 | 80 | 80 | 80 | 80 | 80 | 80 | 80 | 80 | 80 | 80 | 80 |
| **Y** | 75 | 76 | 82 | 83 | 79 | 79 | 79 | 79 | 79 | 83 | 83 | 78 | 86 | 80 | 79 | 79 | 87 | 79 | 79 | 80 | 80 | 99 | 79 | 79 | 100 | 83 | 83 | 79 | 72 | 76 | 79 | 79 | 79 | 79 | 79 | 79 | 79 | 79 | 79 | 79 | 79 |
| **Z** | 76 | 76 | 91 | 90 | 79 | 79 | 79 | 79 | 79 | 90 | 90 | 79 | 82 | 80 | 79 | 79 | 93 | 80 | 79 | 80 | 80 | 83 | 79 | 79 | 83 | 100 | 90 | 79 | 73 | 76 | 79 | 79 | 79 | 79 | 79 | 79 | 79 | 79 | 79 | 79 | 79 |
| **AA** | 76 | 76 | 89 | 89 | 80 | 80 | 80 | 80 | 79 | 89 | 89 | 79 | 83 | 80 | 80 | 80 | 92 | 80 | 79 | 80 | 80 | 83 | 79 | 80 | 83 | 90 | 100 | 80 | 73 | 76 | 79 | 79 | 79 | 79 | 79 | 79 | 79 | 79 | 79 | 79 | 79 |
| **AB** | 75 | 76 | 79 | 79 | 80 | 97 | 97 | 97 | 97 | 79 | 79 | 79 | 79 | 80 | 80 | 80 | 84 | 90 | 80 | 80 | 80 | 79 | 95 | 80 | 79 | 79 | 80 | 100 | 72 | 76 | 97 | 97 | 97 | 97 | 97 | 97 | 97 | 97 | 93 | 93 | 93 |
| **AC** | 72 | 72 | 73 | 73 | 72 | 72 | 72 | 72 | 72 | 73 | 73 | 72 | 72 | 73 | 72 | 73 | 75 | 72 | 73 | 73 | 73 | 72 | 72 | 73 | 72 | 73 | 73 | 72 | 100 | 72 | 72 | 72 | 72 | 72 | 72 | 72 | 72 | 72 | 72 | 72 | 72 |
| **AD** | 78 | 80 | 76 | 76 | 76 | 76 | 76 | 76 | 76 | 76 | 76 | 75 | 76 | 76 | 76 | 76 | 78 | 76 | 76 | 76 | 76 | 76 | 76 | 76 | 76 | 76 | 76 | 76 | 72 | 100 | 76 | 76 | 76 | 76 | 76 | 76 | 76 | 76 | 76 | 76 | 76 |
| **AE** | 76 | 76 | 79 | 79 | 80 | 97 | 100 | 97 | 97 | 79 | 79 | 79 | 79 | 80 | 80 | 80 | 84 | 90 | 80 | 80 | 80 | 79 | 95 | 80 | 79 | 79 | 79 | 97 | 72 | 76 | 100 | 97 | 97 | 97 | 97 | 100 | 97 | 97 | 93 | 93 | 93 |
| **AF** | 76 | 76 | 79 | 79 | 80 | 97 | 97 | 97 | 97 | 79 | 79 | 79 | 79 | 80 | 80 | 80 | 84 | 90 | 80 | 80 | 80 | 79 | 95 | 80 | 79 | 79 | 79 | 97 | 72 | 76 | 97 | 100 | 100 | 100 | 100 | 97 | 100 | 100 | 94 | 94 | 94 |
| **AG** | 76 | 76 | 79 | 79 | 80 | 97 | 97 | 97 | 97 | 79 | 79 | 79 | 79 | 80 | 80 | 80 | 84 | 90 | 80 | 80 | 80 | 79 | 95 | 80 | 79 | 79 | 79 | 97 | 72 | 76 | 97 | 100 | 100 | 100 | 100 | 97 | 100 | 100 | 94 | 94 | 94 |
| **AH** | 76 | 76 | 79 | 79 | 80 | 97 | 97 | 97 | 97 | 79 | 79 | 79 | 79 | 80 | 80 | 80 | 84 | 90 | 80 | 80 | 80 | 79 | 95 | 80 | 79 | 79 | 79 | 97 | 72 | 76 | 97 | 100 | 100 | 100 | 100 | 97 | 100 | 100 | 94 | 94 | 94 |
| **AI** | 76 | 76 | 79 | 79 | 80 | 97 | 97 | 97 | 97 | 79 | 79 | 79 | 79 | 80 | 80 | 80 | 84 | 90 | 80 | 80 | 80 | 79 | 95 | 80 | 79 | 79 | 79 | 97 | 72 | 76 | 97 | 100 | 100 | 100 | 100 | 97 | 100 | 100 | 94 | 94 | 94 |

| | | | | | | | | | | | | | | | | | | | | | | | | | | | | | | | | | | | | | | | | | |
|---|---|---|---|---|---|---|---|---|---|---|---|---|---|---|---|---|---|---|---|---|---|---|---|---|---|---|---|---|---|---|---|---|---|---|---|---|---|---|---|---|---|
| **AJ** | 76 | 76 | 79 | 79 | 80 | 97 | 100 | 97 | 97 | 79 | 79 | 79 | 79 | 80 | 80 | 80 | 84 | 90 | 80 | 80 | 80 | 79 | 95 | 80 | 79 | 79 | 79 | 97 | 72 | 76 | 100 | 97 | 97 | 97 | 97 | 100 | 97 | 97 | 93 | 93 | 93 |
| **AK** | 76 | 76 | 79 | 79 | 80 | 97 | 97 | 97 | 97 | 79 | 79 | 79 | 79 | 80 | 80 | 80 | 84 | 90 | 80 | 80 | 80 | 79 | 95 | 80 | 79 | 79 | 79 | 97 | 72 | 76 | 97 | 100 | 100 | 100 | 100 | 97 | 100 | 100 | 94 | 94 | 94 |
| **AL** | 76 | 76 | 79 | 79 | 80 | 97 | 97 | 97 | 97 | 79 | 79 | 79 | 79 | 80 | 80 | 80 | 84 | 90 | 80 | 80 | 80 | 79 | 95 | 80 | 79 | 79 | 79 | 97 | 72 | 76 | 97 | 100 | 100 | 100 | 100 | 97 | 100 | 100 | 94 | 94 | 94 |
| **AM** | 75 | 76 | 79 | 79 | 80 | 93 | 93 | 93 | 93 | 79 | 79 | 79 | 79 | 80 | 80 | 80 | 84 | 90 | 80 | 80 | 80 | 79 | 93 | 80 | 79 | 79 | 79 | 93 | 72 | 76 | 93 | 94 | 94 | 94 | 94 | 93 | 94 | 94 | 100 | 100 | 100 |
| **AN** | 75 | 76 | 79 | 79 | 80 | 93 | 93 | 93 | 93 | 79 | 79 | 79 | 79 | 80 | 80 | 80 | 84 | 90 | 80 | 80 | 80 | 79 | 93 | 80 | 79 | 79 | 79 | 93 | 72 | 76 | 93 | 94 | 94 | 94 | 94 | 93 | 94 | 94 | 100 | 100 | 100 |
| **AO** | 75 | 76 | 79 | 79 | 80 | 93 | 93 | 93 | 93 | 79 | 79 | 79 | 79 | 80 | 80 | 80 | 84 | 90 | 80 | 80 | 80 | 79 | 93 | 80 | 79 | 79 | 79 | 93 | 72 | 76 | 93 | 94 | 94 | 94 | 94 | 93 | 94 | 94 | 100 | 100 | 100 |

877
