## Supplemental Table 3 for "Lanthanide-dependent isolation of phyllosphere methylotrophs selects for a phylogenetically conserved but metabolically diverse community"

**Table S3.** Significant differences in growth rates and final yields between all strains on 20 mM methanol **(Figure 3) ,** 34 mM ethanol **(Figure 3),** 5 and 12 mM vanillic acid (**Figure 5**), 15 mM methylamine **(Figure 6**)  +/- $LaCl_3$ determined using one-way ANOVA with post-hoc Tukey HSD. Significant differences for each strain +/- $LaCl_3$ or +/- IPTG for lanmodulin expression phenotypes (**Figure 4**) determined by Student's paired t-test

| **20 mM Methanol ± $LaCl_3$** | | | |
|---|---|---|---|
| **Growth Rate** | | **Final Yield** | |
| **Comparison** | **p-value** | **Comparison** | **p-value** |
| SLI 231 ± $LaCl_3$ | 0.005752 | SLI 231 ± $LaCl_3$ | 0.1264 |
| SLI 233 ± $LaCl_3$ | 0.002731 | SLI 233 ± $LaCl_3$ | 0.4189 |
| SLI 384 ± $LaCl_3$ | 0.6598 | SLI 384 ± $LaCl_3$ | 0.6513 |
| SLI 499 ± $LaCl_3$ | 0.00311 | SLI 499 ± $LaCl_3$ | 0.2585 |
| SLI 505 ± $LaCl_3$ | 0.004484 | SLI 505 ± $LaCl_3$ | 0.2158 |
| SLI 575 ± $LaCl_3$ | 0.00002213 | SLI 575 ± $LaCl_3$ | 0.2463 |
| SLI 231 x SLI 233 - $LaCl_3$ | 1 | SLI 231 x SLI 233 - $LaCl_3$ | 0.9232 |
| SLI 231 x SLI 384 - $LaCl_3$ | 0.133 | SLI 231 x SLI 384 - $LaCl_3$ | 0.9546 |
| SLI 231 x SLI 499 - $LaCl_3$ | 1 | SLI 231 x SLI 499 - $LaCl_3$ | 0.9966 |
| SLI 231 x SLI 505 - $LaCl_3$ | 0.9998 | SLI 231 x SLI 505 - $LaCl_3$ | 0.6545 |
| SLI 231 x SLI 575 - $LaCl_3$ | 0.7947 | SLI 231 x SLI 575 - $LaCl_3$ | 0.05092 |
| SLI 233 x SLI 384 - $LaCl_3$ | 0.1335 | SLI 233 x SLI 384 - $LaCl_3$ | 1 |
| SLI 233 x SLI 499 - $LaCl_3$ | 1 | SLI 233 x SLI 499 - $LaCl_3$ | 0.9949 |
| SLI 233 x SLI 505 - $LaCl_3$ | 0.9998 | SLI 233 x SLI 505 - $LaCl_3$ | 0.9861 |
| SLI 233 x SLI 575 - $LaCl_3$ | 0.7929 | SLI 233 x SLI 575 - $LaCl_3$ | 0.01978 |
| SLI 384 x SLI 499 - $LaCl_3$ | 0.1097 | SLI 384 x SLI 499 - $LaCl_3$ | 0.9987 |
| SLI 384 x SLI 505 - $LaCl_3$ | 0.1007 | SLI 384 x SLI 505 - $LaCl_3$ | 0.9702 |
| SLI 384 x SLI 575 - $LaCl_3$ | 0.03403 | SLI 384 x SLI 575 - $LaCl_3$ | 0.02226 |
| SLI 499 x SLI 505 - $LaCl_3$ | 1 | SLI 499 x SLI 505 - $LaCl_3$ | 0.8666 |
| SLI 499 x SLI 575 - $LaCl_3$ | 0.8691 | SLI 499 x SLI 575 - $LaCl_3$ | 0.03202 |
| SLI 505 x SLI 575 - $LaCl_3$ | 0.8975 | SLI 505 x SLI 575 - $LaCl_3$ | 0.01123 |
| SLI 231 x SLI 233 + $LaCl_3$ | 0.5873 | SLI 231 x SLI 233 + $LaCl_3$ | 0.9674 |
| SLI 231 x SLI 384 + $LaCl_3$ | 0.8102 | SLI 231 x SLI 384 + $LaCl_3$ | 0.9996 |
| SLI 231 x SLI 499 + $LaCl_3$ | 0.9683 | SLI 231 x SLI 499 + $LaCl_3$ | 0.9961 |
| SLI 231 x SLI 505 + $LaCl_3$ | 0.6435 | SLI 231 x SLI 505 + $LaCl_3$ | 0.07812 |
| SLI 231 x SLI 575 + $LaCl_3$ | 0.9997 | SLI 231 x SLI 575 + $LaCl_3$ | 0.00001957 |
| SLI 233 x SLI 384 + $LaCl_3$ | 0.9966 | SLI 233 x SLI 384 + $LaCl_3$ | 0.8901 |
| SLI 233 x SLI 499 + $LaCl_3$ | 0.28 | SLI 233 x SLI 499 + $LaCl_3$ | 0.82 |
| SLI 233 x SLI 505 + $LaCl_3$ | 1 | SLI 233 x SLI 505 + $LaCl_3$ | 0.1797 |
| SLI 233 x SLI 575 + $LaCl_3$ | 0.7234 | SLI 233 x SLI 575 + $LaCl_3$ | 0.00001466 |
| SLI 384 x SLI 499 + $LaCl_3$ | 0.4465 | SLI 384 x SLI 499 + $LaCl_3$ | 1 |

| | | | |
|---|---|---|---|
| SLI 384 x SLI 505 + $LaCl_3$ | 0.9991 | SLI 384 x SLI 505 + $LaCl_3$ | 0.05721 |
| SLI 384 x SLI 575 + $LaCl_3$ | 0.9146 | SLI 384 x SLI 575 + $LaCl_3$ | 0.00002191 |
| SLI 499 x SLI 505 + $LaCl_3$ | 0.3154 | SLI 499 x SLI 505 + $LaCl_3$ | 0.04763 |
| SLI 499 x SLI 575 + $LaCl_3$ | 0.8993 | SLI 499 x SLI 575 + $LaCl_3$ | 0.00002345 |
| SLI 505 x SLI 575 + $LaCl_3$ | 0.7776 | SLI 505 x SLI 575 + $LaCl_3$ | 0.00000566 |

| 34 mM Ethanol ± $LaCl_3$ | | | |
|---|---|---|---|
| **Growth Rate** | | **Final Yield** | |
| **Comparison** | **p-value** | **Comparison** | **p-value** |
| SLI 231 ± $LaCl_3$ | 0.09398 | SLI 231 ± $LaCl_3$ | 0.9407 |
| SLI 233 ± $LaCl_3$ | 0.07046 | SLI 233 ± $LaCl_3$ | 0.6408 |
| SLI 384 ± $LaCl_3$ | 0.08337 | SLI 384 ± $LaCl_3$ | 0.8935 |
| SLI 499 ± $LaCl_3$ | 0.3597 | SLI 499 ± $LaCl_3$ | 0.2088 |
| SLI 505 ± $LaCl_3$ | 0.3525 | SLI 505 ± $LaCl_3$ | 0.008165 |
| SLI 575 ± $LaCl_3$ | 0.5879 | SLI 575 ± $LaCl_3$ | 0.694 |
| SLI 231 x SLI 233 - $LaCl_3$ | 0.8923 | SLI 231 x SLI 233 - $LaCl_3$ | 1 |
| SLI 231 x SLI 384 - $LaCl_3$ | 0.1229 | SLI 231 x SLI 384 - $LaCl_3$ | 1 |
| SLI 231 x SLI 499 - $LaCl_3$ | 0.9989 | SLI 231 x SLI 499 - $LaCl_3$ | 0.9171 |
| SLI 231 x SLI 505 - $LaCl_3$ | 0.9319 | SLI 231 x SLI 505 - $LaCl_3$ | 1 |
| SLI 231 x SLI 575 - $LaCl_3$ | 0.2853 | SLI 231 x SLI 575 - $LaCl_3$ | 0.06206 |
| SLI 233 x SLI 384 - $LaCl_3$ | 0.5211 | SLI 233 x SLI 384 - $LaCl_3$ | 0.9999 |
| SLI 233 x SLI 499 - $LaCl_3$ | 0.9805 | SLI 233 x SLI 499 - $LaCl_3$ | 0.9522 |
| SLI 233 x SLI 505 - $LaCl_3$ | 0.4079 | SLI 233 x SLI 505 - $LaCl_3$ | 0.9998 |
| SLI 233 x SLI 575 - $LaCl_3$ | 0.8289 | SLI 233 x SLI 575 - $LaCl_3$ | 0.05244 |
| SLI 384 x SLI 499 - $LaCl_3$ | 0.2151 | SLI 384 x SLI 499 - $LaCl_3$ | 0.9858 |
| SLI 384 x SLI 505 - $LaCl_3$ | 0.02644 | SLI 384 x SLI 505 - $LaCl_3$ | 1 |
| SLI 384 x SLI 575 - $LaCl_3$ | 0.9918 | SLI 384 x SLI 575 - $LaCl_3$ | 0.07454 |
| SLI 499 x SLI 505 - $LaCl_3$ | 0.7844 | SLI 499 x SLI 505 - $LaCl_3$ | 0.9889 |
| SLI 499 x SLI 575 - $LaCl_3$ | 0.4549 | SLI 499 x SLI 575 - $LaCl_3$ | 0.2034 |
| SLI 505 x SLI 575 - $LaCl_3$ | 0.06793 | SLI 505 x SLI 575 - $LaCl_3$ | 0.07879 |
| SLI 231 x SLI 233 + $LaCl_3$ | 0.9987 | SLI 231 x SLI 233 + $LaCl_3$ | 0.9963 |
| SLI 231 x SLI 384 + $LaCl_3$ | 0.004637 | SLI 231 x SLI 384 + $LaCl_3$ | 0.9995 |
| SLI 231 x SLI 499 + $LaCl_3$ | 0.93 | SLI 231 x SLI 499 + $LaCl_3$ | 0.2218 |
| SLI 231 x SLI 505 + $LaCl_3$ | 0.2374 | SLI 231 x SLI 505 + $LaCl_3$ | 0.1215 |
| SLI 231 x SLI 575 + $LaCl_3$ | 0.6082 | SLI 231 x SLI 575 + $LaCl_3$ | 0.111 |
| SLI 233 x SLI 384 + $LaCl_3$ | 0.008673 | SLI 233 x SLI 384 + $LaCl_3$ | 0.9655 |
| SLI 233 x SLI 499 + $LaCl_3$ | 0.7734 | SLI 233 x SLI 499 + $LaCl_3$ | 0.1068 |
| SLI 233 x SLI 505 + $LaCl_3$ | 0.1337 | SLI 233 x SLI 505 + $LaCl_3$ | 0.05601 |
| SLI 233 x SLI 575 + $LaCl_3$ | 0.4013 | SLI 233 x SLI 575 + $LaCl_3$ | 0.05096 |
| SLI 384 x SLI 499 + $LaCl_3$ | 0.001055 | SLI 384 x SLI 499 + $LaCl_3$ | 0.3409 |
| SLI 384 x SLI 505 + $LaCl_3$ | 0.000131 | SLI 384 x SLI 505 + $LaCl_3$ | 0.1958 |
| SLI 384 x SLI 575 + $LaCl_3$ | 0.000385 | SLI 384 x SLI 575 + $LaCl_3$ | 0.1798 |
| SLI 499 x SLI 505 + $LaCl_3$ | 0.7021 | SLI 499 x SLI 505 + $LaCl_3$ | 0.9985 |
| SLI 499 x SLI 575 + $LaCl_3$ | 0.9815 | SLI 499 x SLI 575 + $LaCl_3$ | 0.9971 |
| SLI 505 x SLI 575 + $LaCl_3$ | 0.9678 | SLI 505 x SLI 575 + $LaCl_3$ | 1 |

| 15 mM Methylamine ± $LaCl_3$ | | | |
|---|---|---|---|
| **Growth Rate** | | **Final Yield** | |
| **Comparison** | **p-value** | **Comparison** | **p-value** |
| SLI 231 ± $LaCl_3$ | 0.2622 | SLI 231 ± $LaCl_3$ | 0.8076 |
| SLI 233 ± $LaCl_3$ | 0.3036 | SLI 233 ± $LaCl_3$ | 0.3688 |
| SLI 384 ± $LaCl_3$ | 0.1904 | SLI 384 ± $LaCl_3$ | 0.2502 |
| SLI 499 ± $LaCl_3$ | 0.5452 | SLI 499 ± $LaCl_3$ | 0.6538 |
| SLI 505 ± $LaCl_3$ | 0.8891 | SLI 505 ± $LaCl_3$ | 0.5779 |
| SLI 575 ± $LaCl_3$ | 0.6923 | SLI 575 ± $LaCl_3$ | 0.687 |
| SLI 231 x SLI 233 - $LaCl_3$ | 0.9982 | SLI 231 x SLI 233 - $LaCl_3$ | 1 |
| SLI 231 x SLI 384 - $LaCl_3$ | 0.000869 | SLI 231 x SLI 384 - $LaCl_3$ | 0.00843 |
| SLI 231 x SLI 499 - $LaCl_3$ | 0.8468 | SLI 231 x SLI 499 - $LaCl_3$ | 0.9972 |
| SLI 231 x SLI 505 - $LaCl_3$ | 0.7692 | SLI 231 x SLI 505 - $LaCl_3$ | 0.9984 |
| SLI 231 x SLI 575 - $LaCl_3$ | 0.9989 | SLI 231 x SLI 575 - $LaCl_3$ | 0.02364 |
| SLI 233 x SLI 384 - $LaCl_3$ | 0.001113 | SLI 233 x SLI 384 - $LaCl_3$ | 0.008092 |
| SLI 233 x SLI 499 - $LaCl_3$ | 0.9659 | SLI 233 x SLI 499 - $LaCl_3$ | 0.9984 |
| SLI 233 x SLI 505 - $LaCl_3$ | 0.9256 | SLI 233 x SLI 505 - $LaCl_3$ | 0.9991 |
| SLI 233 x SLI 575 - $LaCl_3$ | 0.9681 | SLI 233 x SLI 575 - $LaCl_3$ | 0.02556 |
| SLI 384 x SLI 499 - $LaCl_3$ | 0.001847 | SLI 384 x SLI 499 - $LaCl_3$ | 0.005875 |
| SLI 384 x SLI 505 - $LaCl_3$ | 0.002082 | SLI 384 x SLI 505 - $LaCl_3$ | 0.00611 |
| SLI 384 x SLI 575 - $LaCl_3$ | 0.0007 | SLI 384 x SLI 575 - $LaCl_3$ | 0.858 |
| SLI 499 x SLI 505 - $LaCl_3$ | 1 | SLI 499 x SLI 505 - $LaCl_3$ | 1 |
| SLI 499 x SLI 575 - $LaCl_3$ | 0.6775 | SLI 499 x SLI 575 - $LaCl_3$ | 0.01568 |
| SLI 505 x SLI 575 - $LaCl_3$ | 0.5903 | SLI 505 x SLI 575 - $LaCl_3$ | 0.0164 |
| SLI 231 x SLI 233 + $LaCl_3$ | 1 | SLI 231 x SLI 233 + $LaCl_3$ | 0.9982 |
| SLI 231 x SLI 384 + $LaCl_3$ | 0.00001482 | SLI 231 x SLI 384 + $LaCl_3$ | 0.0002457 |
| SLI 231 x SLI 499 + $LaCl_3$ | 0.5225 | SLI 231 x SLI 499 + $LaCl_3$ | 1 |
| SLI 231 x SLI 505 + $LaCl_3$ | 0.845 | SLI 231 x SLI 505 + $LaCl_3$ | 0.869 |
| SLI 231 x SLI 575 + $LaCl_3$ | 0.9733 | SLI 231 x SLI 575 + $LaCl_3$ | 0.002645 |
| SLI 233 x SLI 384 + $LaCl_3$ | 0.00001392 | SLI 233 x SLI 384 + $LaCl_3$ | 0.0003005 |
| SLI 233 x SLI 499 + $LaCl_3$ | 0.4466 | SLI 233 x SLI 499 + $LaCl_3$ | 0.9939 |
| SLI 233 x SLI 505 + $LaCl_3$ | 0.7702 | SLI 233 x SLI 505 + $LaCl_3$ | 0.9749 |
| SLI 233 x SLI 575 + $LaCl_3$ | 0.9913 | SLI 233 x SLI 575 + $LaCl_3$ | 0.003543 |
| SLI 384 x SLI 499 + $LaCl_3$ | 0.0000301 | SLI 384 x SLI 499 + $LaCl_3$ | 0.0002317 |
| SLI 384 x SLI 505 + $LaCl_3$ | 0.00002318 | SLI 384 x SLI 505 + $LaCl_3$ | 0.0004394 |
| SLI 384 x SLI 575 + $LaCl_3$ | 0.00001136 | SLI 384 x SLI 575 + $LaCl_3$ | 0.0548 |
| SLI 499 x SLI 505 + $LaCl_3$ | 0.9809 | SLI 499 x SLI 505 + $LaCl_3$ | 0.8198 |
| SLI 499 x SLI 575 + $LaCl_3$ | 0.2506 | SLI 499 x SLI 575 + $LaCl_3$ | 0.002429 |
| SLI 505 x SLI 575 + $LaCl_3$ | 0.4965 | SLI 505 x SLI 575 + $LaCl_3$ | 0.006196 |
| **12 mM Vanillic Acid ± $LaCl_3$** | | | |
| **Growth Rate** | | **Final Yield** | |
| **Comparison** | **p-value** | **Comparison** | **p-value** |
| SLI 231 ± $LaCl_3$ | 0.02098 | SLI 231 ± $LaCl_3$ | 0.1654 |

| | | | |
|---|---|---|---|
| SLI 233 ± $LaCl_3$ | 0.06785 | SLI 233 ± $LaCl_3$ | 0.35 |
| SLI 499 ± $LaCl_3$ | 0.1018 | SLI 499 ± $LaCl_3$ | 0.2796 |
| SLI 505 ± $LaCl_3$ | 0.01559 | SLI 505 ± $LaCl_3$ | 0.01442 |
| SLI 575 ± $LaCl_3$ | 0.8571 | SLI 575 ± $LaCl_3$ | 0.6117 |
| SLI 231 x SLI 233 - $LaCl_3$ | 0.05422 | SLI 231 x SLI 233 - $LaCl_3$ | 0.9758 |
| SLI 231 x SLI 499 - $LaCl_3$ | 0.000385 | SLI 231 x SLI 499 - $LaCl_3$ | 1 |
| SLI 231 x SLI 505 - $LaCl_3$ | 0.1099 | SLI 231 x SLI 505 - $LaCl_3$ | 1 |
| SLI 231 x SLI 575 - $LaCl_3$ | 0.002493 | SLI 231 x SLI 575 - $LaCl_3$ | 0.03123 |
| SLI 233 x SLI 499 - $LaCl_3$ | 0.004892 | SLI 233 x SLI 499 - $LaCl_3$ | 0.9737 |
| SLI 233 x SLI 505 - $LaCl_3$ | 0.8575 | SLI 233 x SLI 505 - $LaCl_3$ | 0.9763 |
| SLI 233 x SLI 575 - $LaCl_3$ | 0.0807 | SLI 233 x SLI 575 - $LaCl_3$ | 0.01756 |
| SLI 499 x SLI 505 - $LaCl_3$ | 0.001474 | SLI 499 x SLI 505 - $LaCl_3$ | 1 |
| SLI 499 x SLI 575 - $LaCl_3$ | 0.1419 | SLI 499 x SLI 575 - $LaCl_3$ | 0.03167 |
| SLI 505 x SLI 575 - $LaCl_3$ | 0.02013 | SLI 505 x SLI 575 - $LaCl_3$ | 0.01953 |
| SLI 231 x SLI 233 + $LaCl_3$ | 0.003649 | SLI 231 x SLI 233 + $LaCl_3$ | 0.5874 |
| SLI 231 x SLI 499 + $LaCl_3$ | 0.05221 | SLI 231 x SLI 499 + $LaCl_3$ | 0.9987 |
| SLI 231 x SLI 505 + $LaCl_3$ | 1 | SLI 231 x SLI 505 + $LaCl_3$ | 0.7883 |
| SLI 231 x SLI 575 + $LaCl_3$ | 0.006383 | 0.SLI 231 x SLI 575 + $LaCl_3$ | 0.0005726 |
| SLI 233 x SLI 499 + $LaCl_3$ | 0.1471 | SLI 233 x SLI 499 + $LaCl_3$ | 0.4589 |
| SLI 233 x SLI 505 + $LaCl_3$ | 0.002128 | SLI 233 x SLI 505 + $LaCl_3$ | 0.98 |
| SLI 233 x SLI 575 + $LaCl_3$ | 0.9563 | SLI 233 x SLI 575 + $LaCl_3$ | 0.0002413 |
| SLI 499 x SLI 505 + $LaCl_3$ | 0.03278 | SLI 499 x SLI 505 + $LaCl_3$ | 0.6414 |
| SLI 499 x SLI 575 + $LaCl_3$ | 0.3161 | SLI 499 x SLI 575 + $LaCl_3$ | 0.0006731 |
| SLI 505 x SLI 575 + $LaCl_3$ | 0.003754 | SLI 505 x SLI 575 + $LaCl_3$ | 0.0001875 |

| **5 mM Vanillic Acid ± $LaCl_3$** | | | |
|---|---|---|---|
| **Growth Rate** | | **Final Yield** | |
| **Comparison** | **p-value** | **Comparison** | **p-value** |
| SLI 231 ± $LaCl_3$ | 0.1459 | SLI 231 ± $LaCl_3$ | 0.0005 |
| SLI 233 ± $LaCl_3$ | 0.7953 | SLI 233 ± $LaCl_3$ | 0.002389 |
| SLI 499 ± $LaCl_3$ | 0.1194 | SLI 499 ± $LaCl_3$ | 0.04408 |
| SLI 505 ± $LaCl_3$ | 0.01286 | SLI 505 ± $LaCl_3$ | 0.02268 |
| SLI 575 ± $LaCl_3$ | 0.6059 | SLI 575 ± $LaCl_3$ | 0.7737 |
| sp. AMS5 ± $LaCl_3$ | 0.08744 | sp. AMS5 ± $LaCl_3$ | 0.4056 |
| SLI 231 x SLI 233 - $LaCl_3$ | 0.788 | SLI 231 x SLI 233 - $LaCl_3$ | 1 |
| SLI 231 x SLI 499 - $LaCl_3$ | 0.9988 | SLI 231 x SLI 499 - $LaCl_3$ | 0.9978 |
| SLI 231 x SLI 505 - $LaCl_3$ | 0.6593 | SLI 231 x SLI 505 - $LaCl_3$ | 0.9547 |
| SLI 231 x SLI 575 - $LaCl_3$ | 1 | SLI 231 x SLI 575 - $LaCl_3$ | 0.000497 |
| SLI 231 x sp. AMS5 - $LaCl_3$ | 0.001096 | SLI 231 x sp. AMS5 - $LaCl_3$ | 0.02093 |
| SLI 233 x SLI 499- $LaCl_3$ | 0.5811 | SLI 233 x SLI 499- $LaCl_3$ | 0.9905 |
| SLI 233 x SLI 505- $LaCl_3$ | 0.1241 | SLI 233 x SLI 505- $LaCl_3$ | 0.9811 |
| SLI 233 x SLI 575- $LaCl_3$ | 0.727 | SLI 233 x SLI 575- $LaCl_3$ | 0.000395 |
| SLI 233 x sp. AMS5 - $LaCl_3$ | 0.000169 | SLI 233 x sp. AMS5 - $LaCl_3$ | 0.01604 |
| SLI 499 x SLI 505- $LaCl_3$ | 0.8523 | SLI 499 x SLI 505- $LaCl_3$ | 0.8003 |

| | | | |
|---|---|---|---|
| SLI 499 x SLI 575 - $LaCl_3$ | 0.9998 | SLI 499 x SLI 575 - $LaCl_3$ | 0.000938 |
| SLI 499 x sp. AMS5- $LaCl_3$ | 0.001957 | SLI 499 x sp. AMS5- $LaCl_3$ | 0.0428 |
| SLI 505 x SLI 575 - $LaCl_3$ | 0.7242 | SLI 505 x SLI 575 - $LaCl_3$ | 0.000151 |
| SLI 505 x sp. AMS5 - $LaCl_3$ | 0.0128 | SLI 505 x sp. AMS5 - $LaCl_3$ | 0.005178 |
| SLI 575 x sp. AMS5- $LaCl_3$ | 0.001309 | SLI 575 x sp. AMS5- $LaCl_3$ | 0.2517 |
| SLI 231 x SLI 233 + $LaCl_3$ | 0.8618 | SLI 231 x SLI 233 + $LaCl_3$ | |
| SLI 231 x SLI 499 + $LaCl_3$ | 0.9976 | SLI 231 x SLI 499 + $LaCl_3$ | 0.9983 |
| SLI 231 x SLI 505 + $LaCl_3$ | 0.04592 | SLI 231 x SLI 505 + $LaCl_3$ | 0.8716 |
| SLI 231 x SLI 575 + $LaCl_3$ | 0.9691 | SLI 231 x SLI 575 + $LaCl_3$ | 0.0002235 |
| SLI 231 x sp. AMS5 + $LaCl_3$ | 3.07E-08 | SLI 231 x sp. AMS5 + $LaCl_3$ | 4.54E-07 |
| SLI 233 x SLI 499 + $LaCl_3$ | 0.6388 | SLI 233 x SLI 499 + $LaCl_3$ | 0.000003121 |
| SLI 233 x SLI 505 + $LaCl_3$ | 0.2714 | SLI 233 x SLI 505 + $LaCl_3$ | 0.6685 |
| SLI 233 x SLI 575+ $LaCl_3$ | 0.9989 | SLI 233 x SLI 575+ $LaCl_3$ | 1.28E-04 |
| SLI 233 x sp. AMS5 + $LaCl_3$ | 1.42E-08 | SLI 233 x sp. AMS5 + $LaCl_3$ | 3.16E-07 |
| SLI 499 x SLI 505 + $LaCl_3$ | 0.02222 | SLI 499 x SLI 505 + $LaCl_3$ | 2.06E-06 |
| SLI 499 x SLI 575 + $LaCl_3$ | 0.8327 | SLI 499 x SLI 575 + $LaCl_3$ | 0.001131 |
| SLI 499 x sp. AMS5 + $LaCl_3$ | 4.21E-08 | SLI 499 x sp. AMS5 + $LaCl_3$ | 0.000001286 |
| SLI 505 x SLI 575 + $LaCl_3$ | 0.1584 | SLI 505 x SLI 575 + $LaCl_3$ | 0.00001043 |
| SLI 505 x sp. AMS5 + $LaCl_3$ | 3.60E-09 | SLI 505 x sp. AMS5 + $LaCl_3$ | 0.001483 |
| SLI 575 x sp. AMS5 + $LaCl_3$ | 1.81E-08 | SLI 575 x sp. AMS5 + $LaCl_3$ | 0.03846 |
| **SLI 575/*lanM* ± IPTG** | | | |
| **Growth Rate in 20 mM MeOH** | | **Growth Rate in 34 mM EtOH** | |
| **Comparison** | **p-value** | **Comparison** | **p-value** |
| 2 μM $LaCl_3$ ± IPTG | 0.9924 | 2 μM $LaCl_3$ ± IPTG | 0.2604 |
| 50 nM $LaCl_3$ ± IPTG | 0.02077 | 50 nM $LaCl_3$ ± IPTG | 0.00736 |
| 1 μM $La_2O_3$ ± IPTG | 0.00772 | 1 μM $La_2O_3$ ± IPTG | 0.000704 |
| 25 nM $La_2O_3$ ± IPTG | 0.01731 | 25 nM $La_2O_3$ ± IPTG | 0.09511 |
